# Precise regulation of the relative rates of surface area and volume synthesis in dynamic environments

**DOI:** 10.1101/806885

**Authors:** Handuo Shi, Yan Hu, Kerwyn Casey Huang

**Affiliations:** Department of Bioengineering, Stanford University, Stanford, CA 94305, USA; Department of Microbiology and Immunology, Stanford University School of Medicine, Stanford, CA 94305, USA; Chan Zuckerberg Biohub, San Francisco, CA 94158

**Keywords:** *Escherichia coli*, *Bacillus subtilis*, *Caulobacter crescentus*, *Schizosacccharomyces pombe*, cell-size control, surface area to volume ratio, cell morphology, cell elongation, cell division

## Abstract

Bacterial cells constantly face complex environmental changes in their natural habitats. While steady-state cell size correlates with nutrient-determined growth rate, it remains unclear how cells regulate their morphology during rapid environmental changes. Here, we systematically quantified cellular dimensions throughout passage cycles of stationary-phase cells diluted into fresh medium and grown back to saturation, and found that cells exhibit characteristic dynamics in surface area to volume ratio (SA/V). SA/V dynamics were conserved across many genetic/chemical perturbations, as well as across species and growth temperatures. We developed a model with a single fitting parameter, the time delay between surface and volume synthesis, that quantitatively explained our SA/V observations, and showed that the time delay was indeed due to differential expression of volume and surface-related genes. The first division after dilution occurred at a tightly controlled SA/V, a previously unrecognized size-control mechanism highlighting the relevance of SA/V. Finally, our time-delay model successfully predicted the quantitative changes in SA/V dynamics due to altered surface area synthesis rates or time delays from translation inhibition. Our minimal model thus provides insight into how cells regulate their morphologies through differential regulation of surface area and volume synthesis and potentiates deep understanding of the connections between growth rate and cell shape in complex environments.

## Introduction

In their natural habitats, bacterial cells constantly face dynamic environmental conditions. To survive, cells alter their physiology to cope with stresses such as nutrient depletion, chemical inhibition, and temperature shifts. During stressful conditions, cells alter their gene expression profiles, often slowing down growth and proliferation and instead allocating limited resources to genes critical for survival^1^. While certain genetic perturbations do not have observable effects in fast-growing cells, they cause death in stressed conditions^2^ or impair survivability when cells resume growth after the environment becomes favorable again^3,4^, highlighting the unique physiological challenges posed by dynamic environments.

Cell shape is intrinsically linked to physiology. During steady-state growth, fast-growing cells in nutrient-rich media adopt larger volumes compared to isogenic cells in minimal media^5^, and systematic tuning of growth rate via medium composition dictates steady-state cell size^6^. Gene expression is also modulated by steady-state growth rates: faster-growing cells tend to have a higher fraction of their proteome devoted to ribosomes, addressing the need for rapid protein synthesis^7,8^. In a batch culture, cell shape can undergo transitions in both cell width and length within several minutes^9,10^, in part as cells adapt their transcriptional program to the new medium and also because the growth of cells can alter the medium composition through nutrient depletion and waste production. Previously, a top-down flux-balance model accurately depicted the kinetics of gene expression and growth in *Escherichia coli* cells under nutrient shifts^11^, but it remains unclear how these environmental and gene expression changes are transduced into cell-shape changes, and whether cell shape is actively optimized in a dynamic environment or is simply a passive outcome of cellular physiology.

In most bacteria, cell shape and size are dictated by the cell wall, a rigid network of peptidoglycan^12,13^. To grow and divide, cells synthesize new peptidoglycan precursors in the cytoplasm, which are then transported to the periplasm and inserted to the expanding cell wall^13^. In rod-shaped bacteria such as *Escherichia coli*, cell elongation and division are regulated by distinct machineries. The actin homolog MreB dictates the insertion pattern of new peptidoglycan material along the cylindrical cell body^14^, which elongates the cell and maintains steady-state cell width^15^. Cell division is regulated by FtsZ, a tubulin homolog that localizes to the mid-cell and forms a ring-like structure prior to division, which then constricts and guides septum formation^13^. Chemical or genetic perturbations to the elongation or division machinery alter cell-shape homeostasis through modified patterns of cell wall synthesis^9,10,15-18^. The extent to which such perturbations disrupt the ability of bacterial cells to adjust to new environments could provide insight into the cellular processes key to shape adaptation.

While cell width and length are thought to be regulated by distinct molecular mechanisms, previous studies have also indicated that they are somewhat inter-connected. In a non-essential gene knockout library, the mutants exhibited variation in both mean cell width and length, with a positive correlation between width and length^19^. Similarly, single point mutations in the MreB protein can alter both width and length^20,21^, yet a large library of mutants were found to all occupy a specific region of the space of cell geometries during growth in LB in which both wider and thinner mutants had longer mean lengths compared to wildtype^10^. Therefore, cell width and length seem to be regulated by an upstream process which unifies the two aspects of cell shape. In a previous study, it was shown that the regulation of surface area to volume ratio (SA/V) is such a process upstream of cell width and length determination: switching cells at steady-state to a condition in which only cell wall (i.e. surface area) synthesis is partially inhibited increased both cell width and length, which lowers the SA/V^22^. Similarly, SA/V changes during the different stages of growth, with log-phase cells having lower SA/V compared to those in stationary phase^22^; those measurements were performed in highly controlled microfluidic chambers, and cells took tens of minutes to several hours to fully adapt to the new steady-state morphology after the almost instantaneous medium switch. The more continuous changes that cells undergo in a dynamic environment such as batch culture have yet to be fully understood, particularly from the perspective of cellular dimensions.

In this study, based on precise and frequent experimental measurements of cellular dimensions and growth rates of a batch culture constantly experiencing nutrient compositional changes and waste accumulations, we develop a model that quantitatively predicts SA/V dynamics. Our model predicts a time delay between surface area and volume synthesis adaptation, and that cells outgrowing from stationary phase will always experience a period of active width increase due to optimal resource allocation to volumetric growth. This model focuses on global resource constraints rather than specific molecular machineries, and therefore is broadly applicable to other microbial batch cultures. Indeed, we found that the observed SA/V dynamics are qualitatively universal across microbial species and growth conditions. With only a single free parameter, our time-delay model predicts the SA/V changes due to perturbations in cell-wall synthesis or protein translation. Our work highlights the ability of bacterial cells to rapidly respond to changing environments by modifying their physical growth.

## Results

### A time-delay model explains the relative dynamics of surface area and volume synthesis in batch cultures

In previous studies, we showed that as *Escherichia coli* cells transition from stationary phase to log phase and back to stationary phase in a batch culture, cellular dimensions vary along with the instantaneous growth rate^9,10^. After a 1:200 back-dilution of an overnight culture grown in LB into fresh LB, cells resumed growth and reached their maximum growth rate after ∼1.5 h, after which growth rate gradually slowed down to approximately zero by ∼4-5 h (Figure 1A). To validate our previous measurements, we extracted a small sample of cells every 15 min and quickly spotted them onto agarose pads for single-cell imaging and quantification (Methods). Mean cell length increased only slightly in the first 0.5 h. The peak in bulk growth rate at 1.5 h corresponded with the peak in mean cell length across the population (Figure 1B), which increased by ∼3-fold relative to stationary phase cells. The mean cell width increased linearly immediately after dilution, and reached its maximum after ∼1 h, increasing by ∼25% relative to stationary phase cells (Figure 1B). Since both length and width initially increase, the surface area-to-volume ratio (SA/V) decreased over this time; SA/V reached its minimum at approximately the same time as the peak in growth rate and mean length (1.5 h; Figure 1C). After 1.5 h, the dynamics of length, width, and SA/V were more gradual, with all quantities reaching plateaus by 5 h. We term these measurements of cell dimensions throughout a passage cycle as a “shape growth curve,” by analogy to absorbance measurements. Using such measurements, we can accurately capture single-cell shape dynamics in a liquid batch culture over extended time periods.

**Figure 1:**
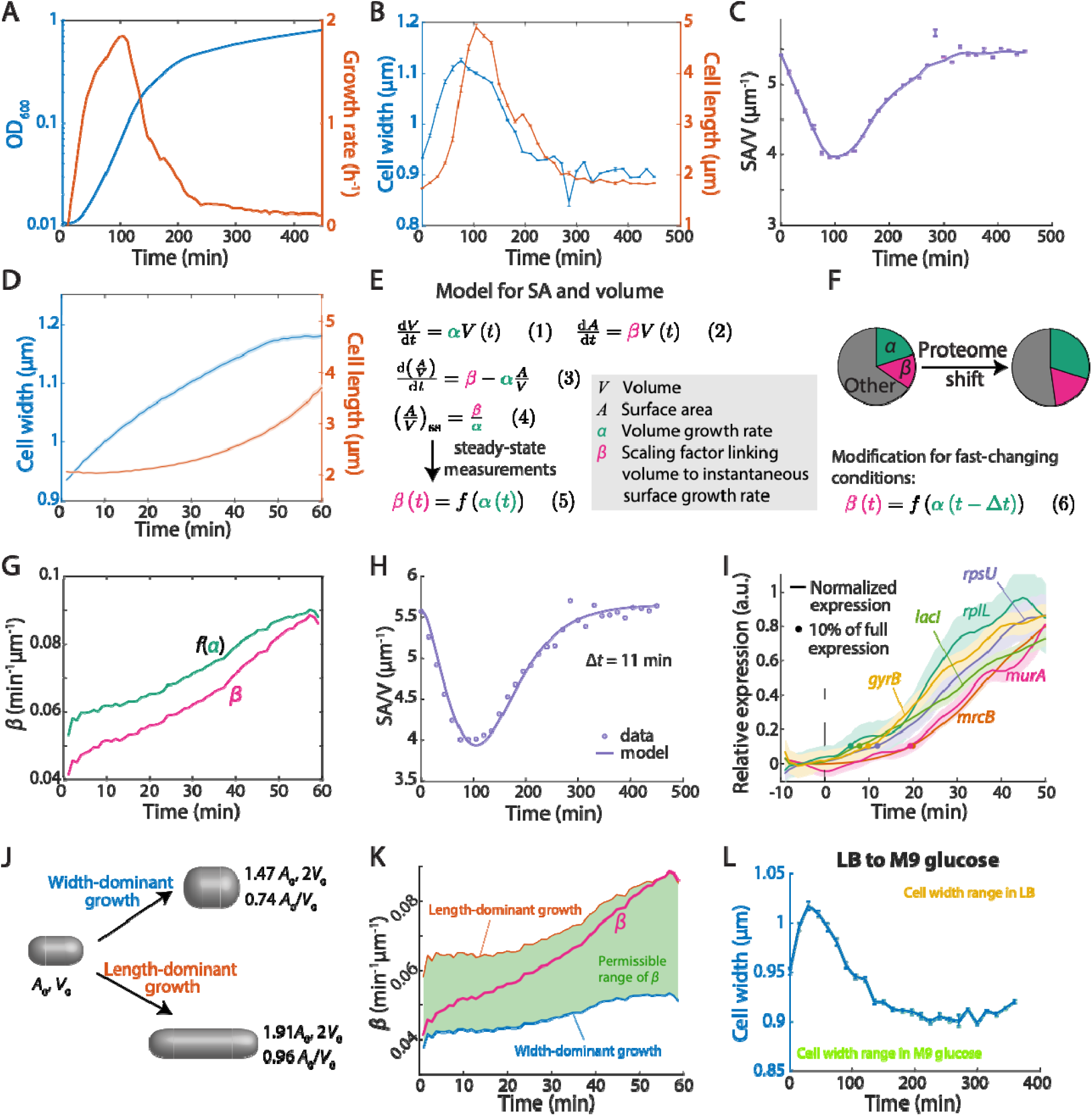
A time-delay model explains the relative dynamics of surface area and volume synthesis in batch cultures. A) Growth curve and the corresponding growth rate of an *E. coli* MG1655 batch culture after diluting the overnight culture 1:200 into fresh LB. Growth rate peaks at ∼100 min post-dilution, and then slowly decreases. B) Shape growth curves (cell width and length as a function of time) for the same culture in (A). Cell width starts to increase earlier compared to length. Data points are mean ± standard error of mean (s.e.m.) for *n* > 200 single cells. C) SA/V as a function of time for the same culture in (B). SA/V decreased as cells resume growth, and then slowly increased back to the initial value when cell growth slowed down. Data points are mean ± s.e.m. with *n* > 200 single cells, and the line is smoothed as a guide to the eye. D) Single-cell time-lapse imaging of stationary-phase MG1655 cells diluted onto an agarose pad containing fresh LB. Each cell exhibited similar width and length dynamics as in the bulk culture in (B). Solid lines and the corresponding shaded areas are mean ± s.e.m. with *n* > 100 single cells. E) A conceptual model for SA/V regulation in cells. The synthesis rates of both surface and volume scale with current cell volume (Eq. 1,2), predicting that the time derivative of A/V depends on *α, β*, and the current A/V (Eq. 3). Steady state A/V is therefore dictated by the ratio between *β* and *α* (Eq. 4). Using previous experimental steady-state SA/V measurements, *β* can be parameterized as a function of *α* in steady state (Eq. 5). F) Given that changes in growth rate (*α*) are accompanied by shifts in proteome composition, our model assumes that the shift is more heavily weighted toward cytoplasmic components than surface area components, thereby causing a delayed change in *β*, which can be approximated by introducing a constant time delay in the function relating *β* to *α* (Eq. 6). G) From the single-cell data in (D), the experimentally measured *β* exhibited a delay of approximately 10 min compared to the steady-state *β* determined by Eq. 5 (i.e. *f*(*α*)). H) Fitting our time-delay model to the experimental data in (C) with Δ*t* = 11 min yielded an excellent fit. I) Protein expression levels were measured using GFP fused to the respective promoters and normalized from 0 to 1. The cytoplasmic proteins increased in expression ∼10 min earlier than the cell-wall synthesis proteins. Dots represent the time points at which expression had increased by 10%. Data are mean ± standard deviation (S.D.) with *n* > 100 cells. J) For a rod-shaped cell starting with surface area *A*_0_ and *V*_0_, doubling its volume by expansion in width costs 47% increase in surface area (top), whereas expansion in length requires a 91% increase in surface area (bottom). Therefore, an increase in width minimizes the surface area requirement for a given amount of volumetric growth. K) Given the geometry and instantaneous growth rate (*α*) of cells in (D), to maintain rod-like shapes, only a certain range of *β* values are permissible (green; Methods). The actual value of *β* started close to its minimal possible value, characterizing a widening-dominant growth mode; by 60 min, *β* reached its maximum possible value, transitioning to an elongation-dominant growth mode. L) Dilution of stationary-phase cells grown in LB into M9 glucose caused increased cell width, despite the fact that cells continuously passaged in M9 glucose always had lower cell width than the LB-grown stationary-phase cells. Data are mean ± s.e.m. with *n* > 100 cells.

To confirm that the observed changes in cellular dimensions and SA/V were not due to artifacts of sampling, we performed time-lapse imaging by diluting and spotting stationary phase cells onto agarose pads containing fresh medium, and tracked the same cells for 1.5 h as they resumed growth on pads. The growth rates of these cells mimicked a population grown in liquid culture, and their shape dynamics recapitulated the shape growth curves (Figure 1D). In particular, each single cell immediately increased in width as soon as it was placed on the agarose pad with fresh medium, while length did not increase noticeably until 20-30 min later (Figure 1D). Therefore, our shape growth curves indeed reflect the morphological changes for each single cell in the batch culture.

Previous work showed that if surface area synthesis, like volume synthesis, is dependent on cell volume (rather than surface area), SA/V will equilibrate at a steady-state value corresponding to the ratio of surface and volume synthesis rates^22^ (Figure 1E). Our measurements for cells transitioning out of and into stationary phase are clearly not at steady state as growth rate is constantly changing, and indeed mean SA/V varied by ∼25%. If we extend the exponential growth laws^22^ for volume 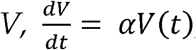, and surface area 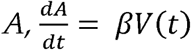, to now cover non-exponential growth through time-dependent functions *α*(*t*) and *β*(*t*), we can derive (Figure 1E) an equation for the dynamics of *A*/*V*:

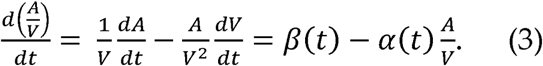

At steady-state, 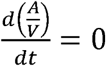, and therefore 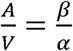. Previous studies have measured these steady-state SA/V values across media that support different growth rates *α*, showing that steady-state SA/V was approximately linear (with negative slope) as a function of *α*^6^. Hence, we approximated *β* as a hyperbolic function of *α* based on our steady-state SA/V measurements (Methods), providing an empirical relationship *β*(*t*) = *f(α*(*t*)).

With our measured value of initial SA/V at *t*=0, the dynamics of *α* (calculated from optical density readouts), and the function *f*, we obtained a prediction for the SA/V dynamics during a shape growth curve (Figure S1A), and found that the model poorly predicted our experimental measurements. Specifically, the model predicted a slower initial decrease in SA/V, and the minimum value was higher and occurred at a later time than our experimental measurements. The final SA/V value never recovered to the initial value, even though the growth rate *α* was 0 at the beginning and end. Thus, the model based on a quasi-steady state assumption does not capture some key factor(s) contributing to the SA/V changes during batch culturing.

Studies focused on the *E. coli* proteome have shown that conditions that support higher growth rate require a reallocation of protein synthesis toward ribosomes^7,8^. We hypothesized that the transition from stationary phase to log growth would require a similar shift in proteome composition more heavily weighted toward cytoplasmic components than surface area components (Figure 1F), which could lead to different temporal dynamics between *α* and *β* contrary to the quasi-steady state hypothesis. To explore such a possibility, we analyzed our single-cell time-lapse trajectories (Figure 1D) to measure *α* and *β* for each cell. From our measurements of *α*, we calculated *f*(*α*), the expected value of *β* during steady-state growth at rate *α*. We found that *f*(*α*) increased more quickly than *β*, with a roughly constant time delay of ∼10 min between the two curves (Figure 1G). Thus, we modified our model in Eq. 3 by substituting the function between *α* and *β* to be *β*(*t*) = *f*(*α*(*t* − *Δt*)), where Δ*t* is a constant that characterizes the time delay between *β* and *α*. Fitting our experimental data with the time-delay model yielded almost perfect agreement with a time delay Δ*t* = 11 min (Figure 1H). Thus, the minimal time-delay model with a single free parameter is able to almost entirely recapitulate the quantitative features of SA/V dynamics.

To confirm that such a delay actually occurs, we utilized a library of *E. coli* strains with GFP reporting the expression from various promoters^23^ and sought to directly quantify the dynamics of cytoplasmic and surface-related proteins as cells emerge from stationary phase. We tested two strains representing ribosomal proteins (P_*rplL*_-GFP, P_*rpsU*_-GFP), two representing other cytoplasmic proteins (P_*lacI*_-GFP, P_*gyrB*_-GFP), and two representing enzymes related to cell-wall synthesis (P_*mrcB*_-GFP, P_*murA*_-GFP). We grew each strain in LB overnight to saturation, then placed the cells directly into a microfluidic device surrounded by the supernatant from the overnight culture. We then switched the spent medium to fresh LB and monitored cell morphology and gene expression. In all six strains, GFP levels increased during growth, signifying increased expression. Consistent with our model prediction, the promoters of cytoplasmic proteins increased expression faster than those of the cell-wall genes (Figure 1I). We quantified the expression dynamics by calculating *t*_10_, the time for each promoter to increase their relative expression by 10% (a proxy for promoter activation). The promoters for cytoplasmic proteins had *t*_10_ ∼ 10 min, whereas the promoters for cell-wall enzymes had *t*_10_ ∼ 20 min, a delay of Δ*t* ∼ 10 min. Therefore, our direct measurement of gene expression dynamics confirmed a delay between *α* and *β*, indicating differential regulation of the proteome (Figure 1F).

Our ability to fit the complex SA/V dynamics during batch culture with a simple model involving only the introduction of a time delay to the steady-state relationship between *β* and *α* suggests a simple picture of the initial stages of growth: as cells emerge from stationary phase, they devote more resources to synthesizing cytoplasmic components such as ribosomes than to surface components such as the cell wall. While the cell still must expand to allow space for these new cytoplasmic components, it does so with a minimal amount of surface growth by expanding predominantly in width rather than length, as volume scales approximately quadratically with width but only linearly with length. Indeed, in a typical rod-shaped cell, a two-fold change in volume requires only a 47% increase in surface area if cells only increase in width, but 91% if cells increase only in length (Figure 1J, Methods). To test this reasoning, from our single-cell time-lapse results (Figure 1D), we directly calculated the possible range of *β* at each time point using the observed width, length and growth rate *α* (Methods). At *t* = 0, the measured *β* was close to its minimal possible value, characterizing a widening-dominant growth mode. By *t* = 60 min, the measured *β* approximately reached its maximum under those growth conditions, consistent with the canonical elongation-dominant growth mode (Figure 1K). Thus, during outgrowth from stationary phase, cells transition their growth mode from widening to elongation.

### Cell widening during exit from stationary phase occurs even if width decreases at steady-state

To further validate that the initial increase in width during outgrowth from stationary phase (Figure 1B,D,K) is governed by the delay between *α* and *β*, rather than determined solely by the current nutrient condition after back dilution, we took stationary-phase cells grown in LB and diluted them into M9 glucose medium. Cells grown in LB always had widths larger than 0.94 µm (Figure 1B), whereas cell widths during passage exclusively in M9 glucose never exceeded 0.94 µm (Figure S1B). Thus, if the medium cells are diluted into dictated cell shape, cell width in M9 glucose would only decrease to below 0.94 µm. However, we observed that by 30 min after dilution into M9 glucose from a stationary-phase LB culture, cell width increased by ∼8% and reached ∼1.03 µm (Figure 1L), a value that was not achievable for cells always passaged in M9 glucose (Figure S1B). Afterward, cell width dropped to ∼0.9 µm, as expected for cells passaged in M9 glucose (Figure S1B), which we presume is mainly dictated by the M9 medium. Therefore, the initial widening of cells during outgrowth is not determined by external nutrients.

Outgrowth from stationary phase also involves changes such as altered expressions of stress-response genes, initiation of DNA replication, and different osmolalities of fresh versus stationary phase media. We tested these possibilities by repeating shape growth curve measurements in a ppGpp^0^ strain and a Δ*thyA* strain. The ppGpp^0^ strain is unable to synthesize ppGpp, a small nucleotide regulating stress-related genes^1^. The Δ*thyA* strain does not replicate DNA in the absence of external thymine^24^. In both strains, shape growth curves still exhibited similar outgrowth dynamics (Supplementary Text, Figure S1C-J). Systematically tuning the osmolality of media caused much smaller shape changes than those observed in the shape growth curves (Supplementary Texts, Figure S1K). Taken together, we conclude that the initial increase in width is likely governed by proteome re-allocation rather than external factors or other biological processes.

### Cells exiting from stationary phase reach a critical SA/V at their first division

Previous work suggested that exponentially growing cells accumulate excess surface area material during the cell cycle, which then triggers division^22^. For a rod-shaped cell, division adds two hemispheric poles and increases SA by 4% without changing volume (Figure S2A). Since cells decrease their SA/V during outgrowth (Figure 1D), we thus asked whether the first division after cells exit from stationary phase was also related to their SA/V.

We diluted stationary-phase cells and placed them onto an agarose pad containing fresh LB, tracked single cells until their first division, and quantified their cellular dimensions. Several models have been proposed for cell-size control during steady-state growth: the “sizer” model, in which cells divide at a fixed size; the “adder” model, in which cells add a fixed volume before division; and the “timer” model, in which cells grow for a fixed time before division^25^. The “sizer” model predicts that the volume at division is independent of initial volume, while the “adder” model predicts a slope of +1 between volume at division and initial volume. During outgrowth from stationary phase, we found that the volume at the first division negatively correlated with the initial volume (Figure 2A), deviating from both the sizer and adder models. Similar negative correlations were observed for cell length and surface area (Figure S2B,C). The timer model also failed to explain the data, with time to first division negatively correlated with initial cell volume (Figure 2B). However, the SA/V at division was approximately constant and independent of the initial SA/V (Figure 2C). Moreover, the normalized distribution of SA/V at division was much narrower compared to other cellular dimensions (Figure 2C,D), suggesting that the first division after stationary-phase exit is linked to a critical SA/V.

**Figure 2:**
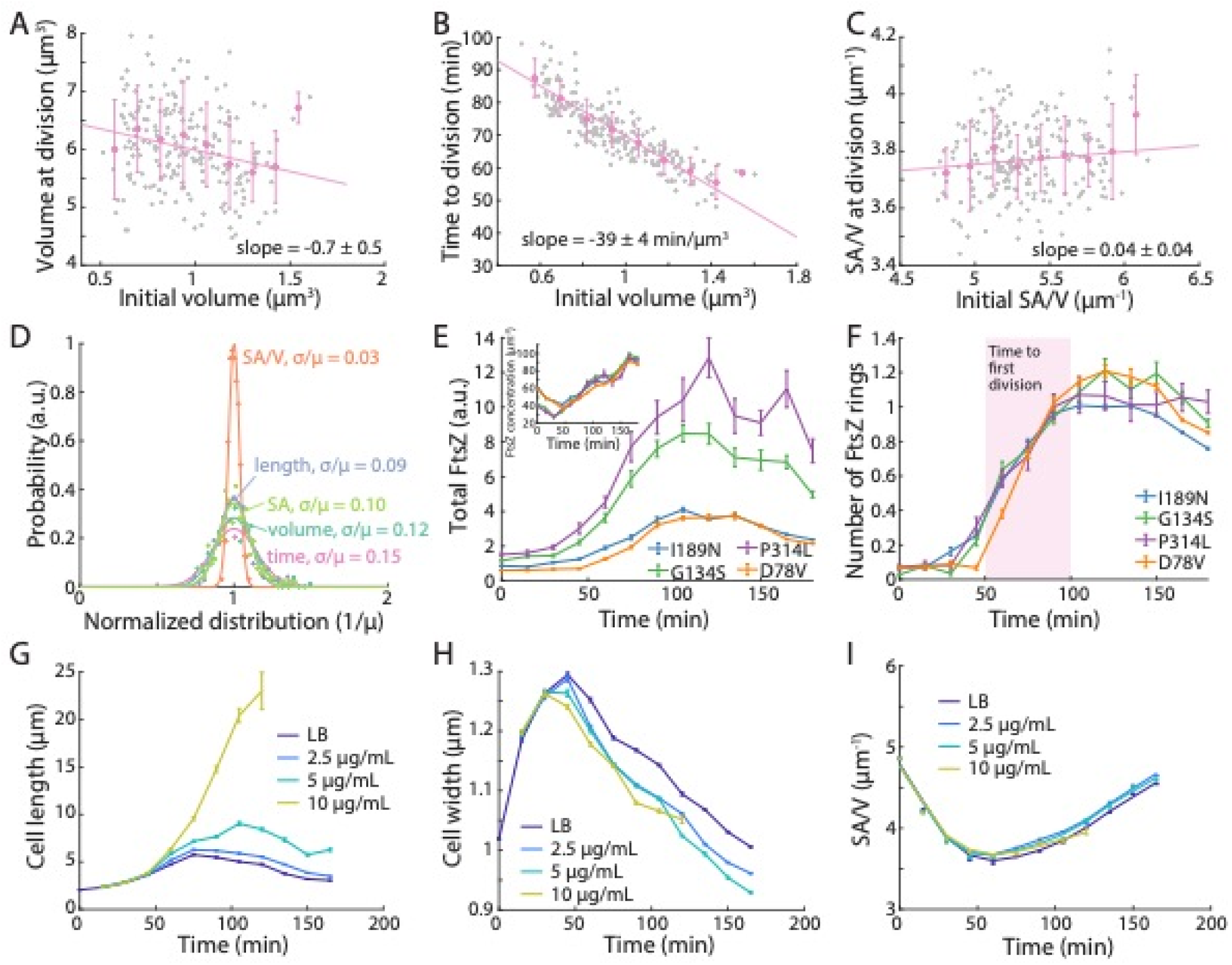
SA/V dynamics are correlated with cell division, and are conserved across cell morphologies. A) For cells exiting stationary phase, the volume at the first division was negatively correlated with the starting volume, deviating from the “sizer” and “adder” models. Gray dots are data points for *n* = 209 individual cells, and pink data points are binned mean and S.D. values, which were fit to a linear model, leading to a slope of −0.7 ± 0.5 (mean ± s.e.m., Pearson’s *r* = −0.22, *p* = 0.001, Student’s *t*-test). B) The time to first division after stationary-phase exit was negatively correlated with the initial cell volume with a slope of −39 ± 4 min/µm^3^ (mean ± s.e.m., Pearson’s *r* = −0.86, *p* < 10^−5^, Student’s *t*-test), indicating that the first division was not regulated by time spent in fresh media. Gray dots are data points for *n* = 209 individual cells, and pink data points are binned mean and S.D. values. C) The SA/V at the first division was largely constant and independent of the initial SA/V of the stationary-phase cell. Gray dots are data points for *n* = 209 individual cells, and pink data points are binned mean and S.D. values, which were fit to a linear model, leading to a slope of −0.04 ± 0.04 (mean ± s.e.m., Pearson’s *r* = 0.11, *p* = 0.12, Student’s *t*-test). D) Normalized distributions of cell length, SA, volume, SA/V, and time at first division after stationary-phase exit. SA/V had by far the narrowest distribution, suggesting that the first cell division after exiting stationary phase occurs precisely at a fixed SA/V. E) FtsZ abundance was measured as total fluorescence intensity inside cells. The dynamics of FtsZ regulation were similar across mutants. FtsZ abundance did not change in the first ∼50 min, then increased to a maximum at ∼100 min. Inset: While larger cells tended to have lower SA/V and higher FtsZ levels, FtsZ concentrations were highly similar across shape mutants. Data points are mean ± s.e.m. with *n* > 200 cells. F) All strains started without FtsZ rings and did not possess a ring until ∼50 min post-dilution, consistent with the onset of first divisions after stationary phase exit (B). By 100 min, virtually all cells had ∼1 FtsZ ring. Data points are mean ± s.e.m. with *n* > 200 cells. G-I) Shape growth curves of *E. coli* MG1655 cells treated with sub-lethal concentrations of cephalexin. Cephalexin causes a dose-dependent increase in cell length (G), accompanied by decreased cell width (H). The increased lengths and decreased widths precisely maintain SA/V for all conditions (I), suggesting that SA/V is robust to perturbations in cellular dimensions. Data points are mean ± s.e.m. with *n* > 200 cells.

We further asked how division-related proteins were regulated during stationary-phase exit by tracking the dynamics of the key division protein FtsZ. We selected four *E. coli* strains, each containing a mutation in the actin homolog MreB that exhibited different mean widths and lengths during log-phase growth ^10^. These mutants allowed us to analyze division dynamics in cells of different widths. These strains also contained a chromosomally-integrated internal fusion of FtsZ to monomeric Venus (FtsZ^sw^-mVenus) at the native FtsZ locus^10,26^, allowing quantification of FtsZ abundance using fluorescence microscopy (Methods). All strains exhibited qualitatively similar cell shape dynamics as wild-type *E. coli*, despite their altered lengths and widths (Figure S2D,E,F). Total FtsZ fluorescence remained constant in the first 45 min post-dilution before starting to increase (Figure 2E), indicating that FtsZ synthesis started much later compared to other genes (Figure 1I). We also measured FtsZ dynamics in a time-lapse experiment with GFP fused to the FtsZ promoter (P_*ftsZ*_-GFP)^23^ and confirmed that FtsZ expression started ∼50 min post-dilution (Figure S2G). Although the four MreB mutants had different levels of total FtsZ (Figure 2E), FtsZ concentration was quantitatively similar across the different strains (Figure 2E, inset), suggesting a conserved mechanism of FtsZ regulation independent of cell shape, consistent with previous measurements in exponential-phase cells^10^. In all strains, no FtsZ rings were observed until ∼50 min post-dilution. Virtually all cells contained one FtsZ ring by ∼100 min post-dilution (Figure 2F), consistent with the observed timing of the first division after stationary-phase exit (Figure 2B). Taken together, these data indicate that FtsZ levels are upregulated concurrent with the need for division.

### SA/V dynamics are broadly conserved across width and length perturbations

Since SA/V is dependent on both cell width and length, we asked whether chemically or genetically tuning cell width or length would affect SA/V dynamics. We first treated wild-type *E. coli* cells with a range of concentrations of cephalexin, a β-lactam antibiotic that inhibits the division-specific cell-wall synthesis enzyme PBP3^27^. Importantly, lower cephalexin concentrations (2.5 µg/mL and 5 µg/mL) did not affect bulk growth rate for at least the first 10 h of growth. For the highest cephalexin concentration used (10 µg/mL), cells started to lyse after 2 h, but growth was not affected prior to lysis (Figure S2H). Thus, cephalexin treatment does not directly affect any parameters in our model, despite the obvious perturbations in cell length. Shape growth curves showed that cells became longer due to inhibition of cell division in a concentration-dependent manner (Figure 2G). However, as cephalexin concentration was increased, cell width peaked at a lower value and began to decrease at an earlier time point (Figure 2H). As a result, SA/V was maintained throughout the first 3 h (Figure 2I) despite the marked changes in cell length. Therefore, cells collectively regulate width and length to maintain SA/V during growth.

We further studied cell shape changes in a library of *E. coli* mutants with a wide range of mean cell lengths and widths^10^. While the strains had different morphologies, they all exhibited similar SA/V dynamics as observed in wild-type *E. coli* (Supplementary Text, Figure S2I-N), further indicating that the SA/V dynamics (Figure 1C) are conserved across genetic perturbations to cellular dimensions.

### SA/V dynamics are conserved across species and growth temperatures in rich media

The basis of our interpretation of SA/V dynamics during batch culture should be generally applicable to species other than *E. coli*, as it does not make any assumptions about *E. coli*-specific pathways. Hence, we predicted that a similar initial decrease in SA/V upon outgrowth from stationary phase should generally occur. We thus expanded our shape growth curve measurements to a variety of other species and conditions. We quantified shape growth curves for the Gram-negative bacteria *Vibrio cholerae* and *Caulobacter crescentus*, both of which form slightly curved rods, the rod-shape Gram-positive bacterium *Bacillus subtilis*, and the eukaryote fission yeast *Schizosaccharomyces pombe*. In all species we tested, SA/V dynamics were similar to those in *E. coli*, with SA/V decreasing when cells resumed growth, and then gradually recovering when cell growth slowed down again (Supplementary Text, Figure S3A-E).

Bacterial cell size during steady-state growth is thought to be relatively constant as growth rate changes across temperatures^5^. Nonetheless, we still found that *E. coli* cells at 30, 37, and 42 °C had different cellular dimensions and SA/V values. At all temperatures, cells still obeyed similar SA/V dynamics (Supplementary Text, Figure S3F,G). Taken together, the initial decreases of SA/V appear to be general across microbial species, growth temperatures, and cell shapes and sizes, indicating that as cells accelerate in growth, volume synthesis always increases more quickly than surface synthesis.

### SA/V regulation is dependent on nutrient conditions

We next asked whether medium composition affects SA/V dynamics, considering that steady-state cell shape is nutrient-dependent^6^. We quantified shape growth curves in M9 media supplemented with different carbon/nitrogen sources that support different growth rates. Overall, in low-nutrient conditions with slower growth rates, Δ*t* was close to zero and SA/V remained largely constant, despite substantial changes in cell width and length (Supplementary Text, Figure S4A-E). Richer nutrient conditions supported faster growth rates and led to larger changes in SA/V and non-zero Δ*t* values (Supplementary Text, Figure S4F-H). Since cell shape changes occurred even in defined media with a single carbon source (Figure S4C,D), the observed cellular dimension changes across shape growth curves are not likely due to diauxic shifts caused by nutrient consumption, but rather related to the dynamics of proteome composition as growth rate changes. A more complex panel of nutrients that support faster growth likely requires a larger shift in proteome composition (Figure 1F), driving a larger range of SA/V changes (Figure S4I).

### Time-delay model quantitatively predicts SA/V decreases due to inhibiting cell-wall synthesis

We next sought to test predictions of our time-delay model experimentally. In our model, reducing *β* (without changing *α*) predicted that the shape growth curve would exhibit larger SA/V decreases during the first 2 h, with a final SA/V when cells returned to stationary phase lower than the initial value (Figure 3A). The *E. coli* cytoplasm is surrounded by the cell envelope, constituted of three layers: the cell wall, and two membranes on either side of the cell wall. The cell wall mainly consists of peptidoglycan, and the membranes are constituted by lipids, proteins, and lipopolysaccharides. Since in most conditions the synthesis of peptidoglycan is more energetically costly compared to lipids^28^, we hypothesized that that altering cell-wall synthesis would have a larger effect on *β*.

**Figure 3:**
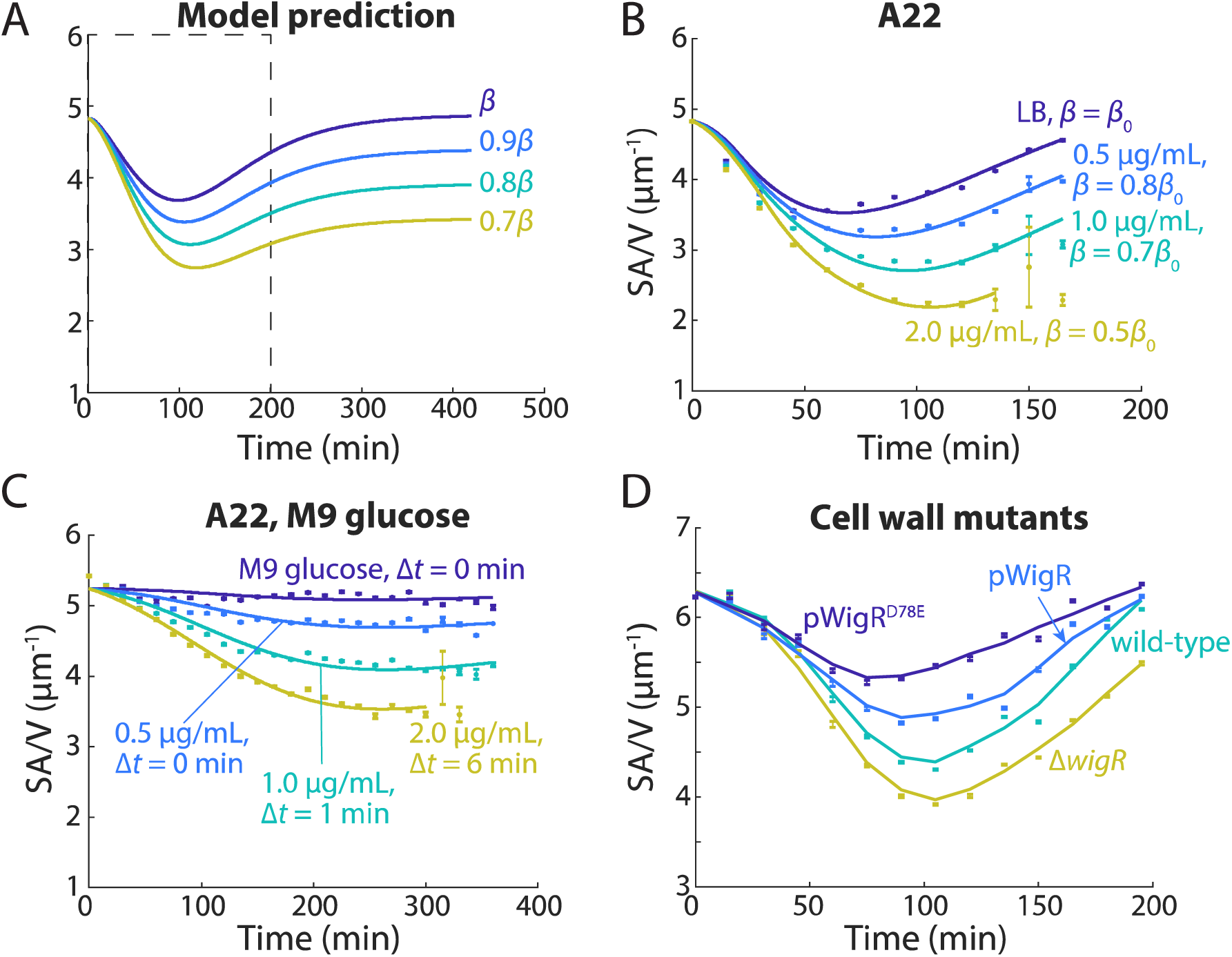
Time-delay model quantitatively predicts SA/V decreases due to inhibiting cell-wall synthesis. A) The model predicts that decreasing *β* results in decreased SA/V over time. B) Shape growth curves during A22 treatment in LB exhibited similar SA/V dynamics as predicted in (A). C) Shape growth curves with A22 treatment in M9 glucose also exhibited decreases in SA/V, even though SA/V remained largely constant without A22. Higher A22 concentrations also led to non-zero Δ*t*, indicating that cells must adjust their proteome under inhibition of cell-wall synthesis. D) Shape growth curves for *V. cholerae* cell-wall synthesis mutants. Overexpression of WigR (pWigR, pWigR^D78E^), which up-regulates cell wall synthesis without affecting growth, increased SA/V. By contrast, Δ*wigR* cells have down-regulated cell-wall synthesis and exhibited lower SA/V.

To validate our model prediction, we treated cells with multiple antibiotics that inhibit cell-wall synthesis and asked whether the SA/V changes were consistent with the model predictions. We first treated MG1655 cells grown in LB with A22^29^, a small molecule inhibiting the actin homolog MreB and therefore affecting the spatial pattern of new cell wall incorporation. Sub-lethal concentrations of A22 did not affect growth rate during the first 3 h of growth, but cells treated with A22 exhibited dose-dependent increases in cell width compared to the non-treated control (Figure S5A), as previously shown ^30^. Interestingly, the increase in cell width was accompanied by relatively lower increases in cell length (Figure S5B), which partially compensated for the decrease in SA/V. Together, the inhibition of cell-wall synthesis by A22 caused SA/V to drop in a dose-dependent fashion (Figure 3B), as predicted by our model (Figure 3A). Fitting the A22 shape growth curves to our model was indeed consistent with a dose-dependent decrease *β* (Figure 3B). Δ*t* was not affected except at the highest A22 concentration, which resulted in a ∼50% longer time delay (Table S2), presumably because higher A22 concentrations require further proteome redistribution. We also tested two other cell-wall synthesis inhibitors, fosfomycin and mecillinam. Fosfomycin inhibits MurA, the enzyme catalyzing the first committed step of peptidoglycan biosynthesis^31,32^, and mecillinam specifically targets PBP2, a transpeptidase that crosslinks new cell-wall material^33^. In both cases, we observed similar changes to shape growth curves as with A22 treatment (Figure S5C,D).

Since cells grown in M9 glucose exhibited largely constant SA/V and virtually no time delay (Figure S4A), we asked whether inhibiting SA synthesis would affect their SA/V dynamics by treating cells grown in M9 glucose with A22. In this case, A22 also caused a similar drop in SA/V (Figure 3C) and mean length (Figure S5E), and cell width now increased as opposed to the slight decrease in the A22-free control (Figure S5F). Thus, the relatively small range of SA/V changes observed in M9 glucose (Figure S4A) was indeed because the proteome composition for surface area and volume synthesis was largely balanced in M9 glucose. Nonetheless, perturbations such as A22 treatment can still break this proteome balance and alter SA/V more substantially. Fitting the A22 SA/V dynamics to our model also showed that at high A22 concentrations, in addition to decreased *β* (Table S2), Δ*t* became non-zero (Figure 3C), suggesting that cells have to re-allocate their proteome composition to accommodate inhibition of cell-wall synthesis.

Cell wall synthesis can also be altered genetically. In *V. cholerae*, activation of a histidine kinase/response regulator two-component system, WigK/WigR, increases expression of many cell-wall synthesis genes and elevates cell-wall synthesis^34^. We therefore quantified the shape growth curves of *V. cholerae* strains with a range of cell-wall synthesis capacities. Compared to wildtype, overexpression of WigR increased SA/V (Figure 3D), mainly through decreased cell width. Overexpression of WigR^D78E^, a phosphomimetic version of WigR, further increases cell wall synthesis^34^, and SA/V increased even more as expected (Figure 3D), accompanied by further decreases in cell width. By contrast, deletion of WigR slows down cell wall synthesis, and we observed decreases in SA/V similar to those during treatment of *E. coli* with wall-acting antibiotics (Figure 3D). Therefore, genetically perturbing cell-wall synthesis affects *β* and therefore SA/V dynamics as predicted by our model.

We next tested whether inhibiting lipid synthesis would have similar effects on SA/V dynamics. While it has been previously shown that inhibiting fatty acid synthesis via treatment with cerulenin alters cell morphology^35^, those experiments were performed in a regime (cerulenin concentration >50 µg/mL) in which growth rates were strongly affected (Figure S5G). In the context of our model, at lower concentrations of cerulenin (<10 µg/mL) where growth rates remained unaffected (Figure S5G), we did not observe noticeable changes in cellular dimensions after 2 h of growth (Figure S5H). Similarly, in cell wall-deficient spheroplasts^36^, surface area was only limited by lipid synthesis, and those cells exhibited increased, rather than decreased, SA/V during outgrowth from stationary phase (Supplementary Text, Figure S5I). We further analyzed previously published proteome datasets^11,37^, and found that levels of lipid synthesis proteins, but not peptidoglycan synthesis proteins, increased monotonically with growth rate (Supplementary Text, Figure S5J-L). Taken together, lipid biosynthesis protein levels are likely directly linked to growth rate, and cell envelope growth is limited by cell wall rather than membrane synthesis.

### Inhibiting translation increases the time delay between volume and surface growth

The addition of the time delay Δ*t* was a critical modification to our model in order to fit our experimental shape growth curve data (Figure 1H, Figure S1A). We therefore asked whether modifying Δ*t* would have observable effects on SA/V dynamics. By increasing Δ*t* from 11 to 25 min, our model predicted that the minimal SA/V reached in log phase would decrease, while the final SA/V when cells enter stationary phase would increase (Figure 4A). Such non-monotonic changes to SA/V dynamics are somewhat counter intuitive, highlighting the biological relevance of the time delay. The initial SA/V drop after exiting stationary phase is primarily due to the quick increase in *α*. While *β* also increases, the time delay between *α* and *β* causes *V* to grow faster than *A*, leading to decreased SA/V. Therefore, a larger Δ*t* means that *V* increases more before *A* growth catches up, resulting in an even lower SA/V in log phase. As for the terminal SA/V, it is dependent on 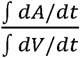, an integral effect that depends on the entirety of the growth dynamics. 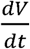 only depends on and remains unaffected, while 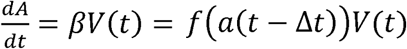. Increasing Δ*t* leads to increased ∫ *dA*/*dt* and eventually a higher terminal SA/V.

**Figure 4:**
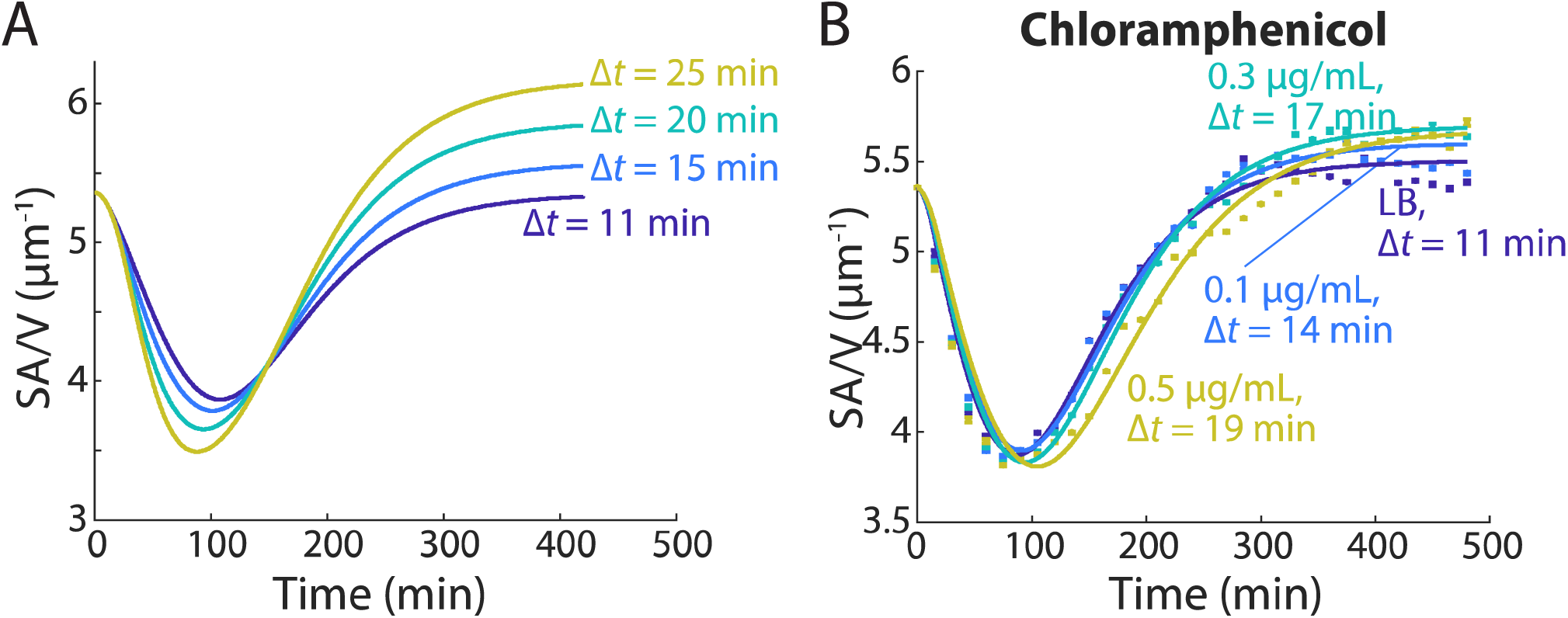
Inhibiting translation increases the time delay between volume and surface growth. A) The model predicts that higher Δ*t* leads to lower SA/V in log phase, but higher SA/V when cells enter stationary phase again. B) Shape growth curves of MG1655 cells treated with low levels of chloramphenicol. The dose-dependent SA/V dynamics were consistent with model predictions, and model fitting resulted in longer Δ*t* at higher doses. Data points are mean ± s.e.m., with *n* > 200. Solid lines are best fits to the time-delay model.

We tested these predictions by treating cells with low levels of chloramphenicol, a translational inhibitor, based on our inference that Δ*t* is mainly determined by rates of proteome re-allocation (Figure 1F). Although inhibiting protein translation inevitably affected both *α* and *β*, at very low chloramphenicol concentrations (∼0.01-0.05X minimal inhibitory concentration), cell growth was largely unaltered, and the SA/V dynamics matched our model predictions: increasing chloramphenicol concentrations increased Δ*t* by ∼70%, while *β* only changed <5% across conditions (Figure 4B). In these cases, the changes in SA/V were largely dictated by the relatively larger cell widths under chloramphenicol treatment (Figure S6A), as cell length followed very similar trends across concentrations (Figure S6B). Higher chloramphenicol concentrations substantially reduced growth rate and led to even lower SA/V in log phase (Figure S6C). Model fitting found that the reduced SA/V was also mainly due to increased *Δt* (Figure S6C), as *β* changed less than 10% (Table S2). Thus, our shape growth curves with chloramphenicol treatment provide further validation of our model predictions, and demonstrate that tuning Δ*t* has observable effects, rather than merely acting as a fitting parameter.

## Discussion

Although much work has been done to unveil the molecular players determining cell morphology at steady states, an approach unifying cell morphology and growth rates in dynamic environments is still lacking. In this study, our time-delay model of SA/V dynamics accurately predicted how cell size responded to growth rate changes, and was generally applicable across many species and growth conditions. The minimal assumption of a single, constant time delay between volume and surface area synthesis quantitatively recapitulated the complex SA/V dynamics (Figure 1C, S3, S4), and predicted SA/V changes under perturbations (Figure 3, 4). While previous studies have shown that cells adopt different steady-state SA/V values across growth conditions^6,22^, our work further reveals that the quantitative dynamics of SA/V under environmental perturbations are based on global resource allocation and the temporal dependence between surface area and volume growth (Figure 1I). Our model thus has the potential to quantitatively describe other dynamic biological processes that scale with cell volume^38^.

While we obtained the time delay between surface area and volume synthesis (*Δt*) via model fitting, systematically tuning *Δt* conferred non-monotonic changes in SA/V dynamics (Figure 4A,B), highlighting its biological relevance. Indeed, we were able to quantify the delay of surface area growth compared to volumetric growth by directly tracking protein expression in single cells (Figure 1G), and the delay was observed previously when cells were switched from one steady state to another^22^. Inhibiting protein translation increased *Δt* (Figure 4B), suggesting that surface area synthesis was delayed due to a slower regulation of surface area synthesis-related enzymes. The connection between *Δt* and changes in growth rate is complex, since higher growth rates may require larger shifts in proteome, but also speeds up proteome turnover. Thus, Δ*t* does not necessarily have a simple correlation with growth rates. Other factors, such as cell-wall precursor synthesis and insertion, could also alter Δ*t* in addition to proteome changes. It remains to be explored why and how surface area and volume synthesis rates are differentially modulated, and the ways in which such a delay can be beneficial for cells.

Our model predicts that upon growth resumption, cells should always increase in width, as surface area synthesis is limiting and increasing width rather than length poses a lower demand for surface area material (Figure 1J). We observed such width increases after diluting stationary-phase cells into fresh media across all species and experimental conditions (Figure 1B,D,L, S3, S4). Similar changes in width have been reported in growth-inhibited cells due to the presence of an antibiotic, in which *E. coli* cells resumed growth after washing out the antibiotic and cell width increased prior to mlength increase^39^. The initial widening of cells is likely related to active growth and the subsequent imbalance between surface area and volume synthesis. In *E. coli*, such widening has been linked to altered cell wall insertion patterns governed by changes in MreB localization patterns^9,10^, providing a potential molecular mechanism in response to limited surface area synthesis. While DNA replication is a central process in cell proliferation and has implications in cell-size determination^6,40^, we have shown that cell shape remodeling can be independent of DNA replication (Figure S1G-J), at least in the initial 2 h of shape growth curves, indicating that the effect of DNA abundance on cell shape potentially manifests on a different time scale or is growth dependent. Regardless, our findings strongly suggest that the dynamic coordination between surface area and volume growth dictates cell shape.

*E. coli* cell division at steady state is well described by the adder model^41^. However, for cells growing out of stationary phase, we observed that cells did not add a constant volume, surface area, or length (Figure 2A, Figure S2B,C), nor did they divide at a fixed size or after a fixed time interval (Figure 2A,B). Instead, cell division occurred at a given SA/V (Figure 2C,D), highlighting the biological relevance of SA/V regulation. The key division protein, FtsZ, was upregulated concurrently with the need for cell division (Figure 2E,F, S2G), consistent with previous work showing that a threshold of FtsZ is required for cell division^42,43^. The dynamics of FtsZ were further delayed compared to other cell-wall synthesis genes (Figure 1I, S2G), reinforcing the idea that gene expression during growth resumption is temporally regulated to allow cells to prioritize volumetric growth over surface area synthesis or division.

Our model of SA/V dynamics links cell width and length, even though the two dimensions are regulated by distinct molecular machineries^14,15,33^. Interestingly, perturbations known to increase cell width or length also led to decreases in the other dimension (Figure 2G,H, S5A,B,E,F). Larger cell width and length both decrease SA/V, and a corresponding decrease in cell length or width can compensate for an increase in the other to maintain SA/V. Therefore, despite the seemingly disjoint nature of cell width and length regulation in rod-shaped cells, the global resource limitation on surface area growth poses constraints on the two dimensions. From an evolutionary perspective, bacterial cells constantly encounter feast or famine conditions, and therefore optimize the resource allocation strategies through regulations between surface area and volume growth. Rod-like shapes allow cells to efficiently tune SA/V via two disparate growth modes (Figure 1J), which potentially confers evolutionary benefits compared to other shapes. In eukaryotic cells, a nuclear transporter receptor serves as a sensor of SA/V^44^, and it remains to be discovered whether similar global SA/V sensors exist in bacterial cells, which would provide opportunities to reveal new connections between cell physiology, size, and fundamental mechanisms of morphogenesis.

## Methods

### Strains and media

Strains used in this study are described in Table S1. For routine culturing, all cells were grown in lysogeny broth (LB) at 37 °C unless otherwise specified. *C. crescentus* cells were grown in PYE (peptone-yeast extract) media at 30 °C as previously described^22^, and *S. pombe* cells were grown in YES255 media at 30 °C. Antibiotics (Sigma Aldrich, St. Louis, MO, USA) were used at the concentrations noted in the text. IPTG was used at a final concentration of 1 mM for *V. cholerae* cells. Thymine was added at a final concentration of 500 µg/mL for the *E. coli* Δ*thyA* strain. For minimal media, glucose or other carbon sources were added at final concentration of 0.4%, six amino acids (L-methionine, L-histidine, L-arginine, L-proline, L-threonine, and L-tryptophan) were added to final concentrations of 500 µg/mL each, and casamino acids were added to a final concentration of 3% (w/v).

Strains were inoculated from freezer stocks into test tubes with 3 mL of media and supplemented with the appropriate antibiotics. The tubes were incubated overnight, except for cells grown in M9 acetate, which were incubated for 48 hours. Overnight cultures were back-diluted 1:200 into the same fresh medium for shape growth curves and growth curves measurements.

Spheroplasts were grown in LFLB (LB with additional 3.6% sucrose and 10 mM MgSO_4_) at 30 °C, with 60 µg/mL cefsulodin added to inhibit cell wall growth^45^. For shape growth curve measurements, overnight spheroplast cultures with cefsulodin were washed three times in fresh LFLB, and diluted 1:10 into LFLB with or without cefsulodin.

### Single-cell imaging

For imaging, samples were taken from test tubes and placed on 1% agarose pads every 15 min, and then imaged within 5 min. For membrane staining, a small aliquot of cells was incubated with FM 4-64 (Invitrogen) at a final concentration of 5 µg/mL for 5 min and spotted on agarose pads without washing. Phase-contrast images and epifluorescence images were acquired with a Nikon Ti-E inverted microscope (Nikon Instruments) using a 100X (NA 1.40) oil immersion objective and a Neo 5.5 sCMOS camera (Andor Technology). The microscope was outfitted with an active-control environmental chamber for temperature regulation (HaisonTech, Taipei, Taiwan). Images were acquired using µManager v.1.4^46^.

### Morphological analyses

The MATLAB (MathWorks, Natick, MA, USA) image processing code *Morphometrics*^19^ was used to segment cells and to identify cell outlines from phase-contrast or fluorescence microscopy images. A local coordinate system was generated for each cell outline using a method adapted from *MicrobeTracker*^47^. Cell widths were calculated by averaging the distances between contour points perpendicular to the cell midline, excluding contour points within the poles and sites of septation. Cell length was calculated as the length of the midline from pole to pole. Cell volume and surface area were estimated from the local meshing results.

### FtsZ fluorescence quantification

Cells with a sandwich fusion of mVenus to FtsZ^26^ were imaged in phase contrast and fluorescence using an ETGFP filter. FtsZ fluorescence was quantified by summing the intensity values of each pixel within the cell contour. FtsZ rings were identified as the peak of fluorescence intensity along the cell contour.

### Population-level growth analyses

To measure growth dynamics, overnight cultures were inoculated into 200 µL of fresh media supplemented with the appropriate antibiotics in a clear 96-well plate. The plate was covered with an optical film, with small holes poked at the side of each well to allow aeration. Incubation and OD measurements were performed with an Epoch 2 plate reader (BioTek) at appropriate temperatures with continuous shaking and OD_600_ measured at 7.5-min intervals. The growth rate was calculated as the slope of ln(OD) with respect to time after smoothing using a moving average filter of window size five.

### *β* as a function of *α* at steady state

Based on previous steady-state growth rate and SA/V measurements, steady-state SA/V linearly correlates with growth rate^6^. Since steady-state SA/V equals *β*/*α*, and growth rate is *α, β*/*α* is a linear function of *α*. Therefore, based on SA/V measurements at two different growth rates, we can fit the function relating *β* to *α*. In our measurements in LB, *E. coli* cells approaching stationary phase correspond to *α* ∼ 0 and *β*/*α* = 5.7 µm^-1^. By diluting cells repeatedly, cells reached steady-state morphologies^10^ for which *β*/*α* = 3.8 µm^-1^, and the corresponding *α* = 1.7 h^-1^. Thus, in LB, we have *β*/*α* = 5.7 – 1.12*α*, or *β* = *α*(5.7 – 1.12*α*).

### SA/V as a function of width and length changes

For calculations in Figure 1J and S2E, a cell with width *w* and length *l* was approximated by a cylindrical cell body with radius *r* = *w*/2 and length *l* – 2*r*, and hemispherical caps on each end with radius *r = w*/2. In this scenario, the corresponding surface area of the cell is *SA = 2πr(l –* 2*r) +* 2 × 2*πr*^2^ *= πwl*, and the volume is *V = πr*^2^*(l –* 2*r) +* 4*πr*^3^/3 = *πw*^2^*l/*4 *– πw*^3^*/*12. For the initial cell, *w =* 1 µm and *l* = 5 µm.

### Calculation of permissible range of *β*

In rod-shaped cells, given the above equations for *V* and *A*,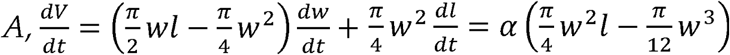,and 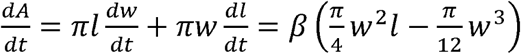. During growth,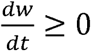 and 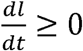. Solving for the linear optimization with the above constraints gives a permissible range of *β* that is dependent on *w, l*, and *α*, with minimum *β* Corresponding to 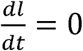, and maximum *β* corresponding to 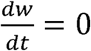.

### DAPI staining and fluorescence quantification

Cells were grown in the same conditions as in single-cell imaging experiments. At each time point, 1 mL of each sample was taken, pelleted at 6,500 rcf for 1 min, and fixed via resuspension in 500 µL 70% ethanol and incubation for 15 min at room temperature. Cells were then pelleted at 6,500 rcf for 1 min, resuspended in 500 µL PBS, with 4’,6-diamidino-2-phenylindole (DAPI) added to a final concentration of 1 µg/mL, and incubated in the dark for 15 min. Cells were washed with PBS twice by pelleting at 6,500 rcf for 1 min followed by resuspension. Cells were spotted onto 1% agarose pads and imaged in phase contrast and fluorescence using a DAPI filter. DAPI fluorescence was quantified by summing the intensity values of each pixel within the cell contour, and then normalizing to the corresponding intensity in the control.

### qPCR

To estimate the relative replication rate of the chromosome, we quantified the copy numbers of 16 genetic loci via qPCR and fitted their log(relative abundance) to their corresponding distances to *terC*, as previously described^18^. The cells were harvested at six different time points (in the overnight culture, and 30, 60, 90, 120, and 150 min after 1:200 dilution), and the DNA was extracted using the genomic DNA purification kit (Qiagen). The relative abundances of chromosomal loci were quantified by qPCR, using the EvaGreen qPCR kit (Bio-rad). The qPCR probes used were as previously described^47^.

## Acknowledgements

The authors thank Alexandre Colavin, Linfeng Yang, Leigh Harris, Po-Yi Ho, and Petra Levin for helpful discussions, and Tobias Dörr and Suckjoon Jun for strains. This work was supported by NIH Director’s New Innovator Awards DP2OD006466 (to K.C.H.), NSF CAREER Award MCB-1149328 (to K.C.H.), the Allen Center for Systems Modeling of Infection (to K.C.H.), and an Agilent Graduate Fellowship and a Stanford Interdisciplinary Graduate Fellowship (to H.S.). K.C.H. is a Chan Zuckerberg Biohub Investigator.

## Data availability statement

The datasets generated during and/or analysed during the current study are available from the corresponding author on reasonable request.

