## Supplementary Materials for "Precise regulation of the relative rates of surface area and volume synthesis in dynamic environments"

**Supplementary Text**

**Cell widening during exit from stationary phase is not dependent on the stringent response, DNA replication, or changes in external medium osmolality**

Nutrient limitation in stationary-phase cells elevates the level of the small nucleotide ppGpp, which then upregulates the expression of stress-related genes^1^. To test whether those stress responses directly caused the changes in SA/V, we measured the shape growth curve of an *E. coli* strain unable to synthesize ppGpp (ppGpp^0^). Although ppGpp^0^ cells were slightly wider and longer than wild-type cells^2^, they still exhibited similar cell shape dynamics, with cell width increasing before length (Figure S1C) and overall lower SA/V during log phase (Figure S1D).

After stationary-phase cells are diluted into fresh medium, multiple cellular processes are activated, including DNA replication, transcription, and translation. Transcription and translation are directly linked to cellular growth and proteome reallocation (Figure 1F), which are accounted for in our model. However, DNA synthesis is somewhat separated from the other processes, hence we wondered whether DNA replication could be upstream of the morphological changes we observed. We quantified the timing of DNA replication via qPCR by measuring the relative abundances of 16 genomic loci spread across the genomes and estimating the progress of replication forks (Methods) for cells grown in LB at different time points after back dilution into fresh medium. Since each round of genomic replication starts at the replication origin (*oriC*) and ends at the terminus (*terC*), during exponential growth, the genomic regions closer to *oriC* will have a higher copy number relative to those closer to *terC*. The rate of DNA replication can therefore be estimated as the slope of log(relative loci abundance) vs. the distance to *terC* ^3^. In stationary phase, DNA replication virtually halts and therefore the relative abundances of all genomic loci should be roughly equal, corresponding to a slope close to zero. Our qPCR results showed that in both the 0-min (i.e. stationary phase) and 30-min cultures, DNA replication rates were close to zero. DNA replication increased after 30 min, approaching the maximum by 90 min (Figure S1E,F). Thus, DNA replication did not start until after cells had already experienced morphological changes at the first 30 min outgrowth, indicating that DNA replication does not control the initial widening of the cells.

We further modulated DNA replication using a strain lacking *thyA*, which encodes the thymidylate synthase in *E. coli*. Δ*thyA* strains require external thymine for growth and DNA replication^4^. We grew Δ*thyA* cells in LB with 500 µg/mL of thymine to stationary phase, and diluted the cells into LB with or without additional thymine. In both cases, bulk growth rates were not affected in the first 4 h of growth. We quantified the relative DNA abundances in both conditions by labeling fixed cells using DAPI (Methods) and verified that absence of thymine indeed slowed down DNA replication in the Δ*thyA* strain (Figure S1G). However, the decreased DNA replication activity did not affect the shape growth curve during the first 2 h of growth (Figure S1H-J), further indicating that the initial widening of the cells after back-dilution is largely independent of DNA replication.

Since cell growth could alter the osmolarity of the medium due to the production or consumption of osmolytes, it was possible that the initial increase in cell width upon dilution into fresh medium was due to a hypoosmotic shock. We directly assayed the osmotic strength of fresh LB and spent LB (the supernatant from a stationary-phase culture grown in LB) using an osmometer. The osmolalities were 252 mOsmol/kg and 262 mOsmol/kg, respectively, indicating that cells underwent a slight 10 mOsmol/kg hypoosmotic shock during the transition. To determine the extent to which such a shock would affect cellular dimensions, we grew *E. coli* cells in LB supplemented with 500 mM of sorbitol (corresponding to an additional 500 mOsmol/kg in osmolality) to stationary phase. We then exposed these cells in a microfluidic device to spent LB with lower concentrations of sorbitol. The stationary-phase cells did not grow due to nutrient limitation, but the osmotic changes caused a hypoosmotic shock and therefore expansion of the cells in both width and length due to water influx that equilibrated within 5-10 min after the media switch. This time scale was short compared to the duration of the observed width increase in shape growth curves, which typically lasted tens of minutes (Figure 1B,L, S1B,C). Moreover, we found that even with an osmolality decrease of 500 mOsmol/kg, cell length increased by only ~7% and width increased by <4% (Figure S1K). Since the actual osmolality differences between fresh vs. spent media were ~50-fold smaller (10 mOsmol/kg), we expect that osmotic effects contribute negligibly to shape growth curves. Moreover, our data demonstrate that osmotic shocks cause larger increases in cell length compared to cell width. Thus, the initial large increase specifically in cell width is highly unlikely to be due to osmotic shock.

**SA/V dynamics are conserved across cell shape mutants**

We previously showed that a library of *E. coli* strains with mutations to the actin homolog MreB in *E. coli* could result in a wide range of mean cell lengths and widths encompassing an ~5-fold range in volume, without affecting maximum growth rates^5^. While it is challenging to quantify the shape growth curves for ~100 strains, we acquired cellular dimensions for stationary phase and log phase (2 h post 1:200 dilution) for all strains to examine whether they followed similar trends as wild-type. Across all strains in the library, mean cell length was higher after 2 h of outgrowth compared with stationary phase, as expected (Figure S2I). Mean cell widths in stationary phase were strongly correlated with widths after 2 h, with most strains exhibiting higher width in log phase (Figure S2J), as for wildtype. However, a minority of strains were wider in stationary phase (Figure S2J) and thus presumably underwent different width dynamics. Nonetheless, SA/V was higher in stationary phase compared with log phase for every strain in the library (Figure S2K), indicating that the SA/V decrease during outgrowth is generic to a wide range of cell sizes. To validate these measurements, we carried out shape growth curves for three mutants with a range of mean cell widths spanning the library; the MreB^E255V^ mutant had lower width in log phase than stationary phase. While the mutants all started with different SA/V values, they followed a similar qualitative trajectory as wild-type cells (Figure S2L). Interestingly, the MreB^E255V^ mutant initially increased in width, but decreased below the stationary phase width value by *t* = 2 h (Figure S2M), consistent with our observations for the mutant library as a whole. While MreB^E255V^ width started to decrease as early as *t* = 1 h, length continued to increase similar to other strains (Figure S2N), counteracting the effects of decreased width on SA/V and thereby maintaining a low and roughly constant SA/V until *t* = 2 h. These data indicate that the typical SA/V dynamics we have described can be achieved despite more complex dynamics in cellular dimensions, and that cell length and width both contribute to SA/V dynamics.

**SA/V dynamics are conserved across species and growth temperatures in rich media**

The basis of our interpretation of SA/V dynamics during batch culture should be generally applicable to species other than *E. coli*, as it does not make any assumptions about *E. coli*-specific pathways. Hence, we predicted that a similar initial decrease in SA/V upon outgrowth from stationary phase should generally occur. We thus expanded our shape growth curve measurements to a variety of other species and conditions. Indeed, the Gram-negative bacterium *Vibrio cholerae*, which forms slightly curved rods, exhibited a SA/V decrease of ~35% over the first 1.5 h post dilution, at which time it reached its maximum growth rate (Figure S3A). The dynamics of cell width and length were similar to those in *E. coli*. We also investigated another Gram-negative bacterium, *Caulobacter crescentus*, which forms a curved rod similar to *V. cholerae* and has a highly regulated cell cycle*.* *C. crescentus* cells maintained mean cell length throughout the first 6 h of measurements. Nonetheless, cell width increased by ~15%, leading to a ~15% decrease in SA/V (Figure S3B).

Unlike those Gram-negative bacteria, the Gram-positive rod *Bacillus subtilis* is generally considered to maintain its width across nutrient conditions. However, we observed an initial ~15% increase in cell width in its shape growth curve, leading to a similar decrease in SA/V (Figure S3C) as in *E. coli*. Since *B. subtilis* cells exhibit chaining during growth^6^, we also used the membrane stain FM 4-64 to quantify cell shape in *B. subtilis*. Despite a quantitative shift in SA/V due to the septa in chaining cells that were not detectable by phase imaging, *B. subtilis* cells exhibited the same qualitative dynamics for cellular dimensions and SA/V (Figure S3D) when measured using membrane staining.

The eukaryote fission yeast *Schizosaccharomyces pombe* is well known to obey sizer regulation during steady-state growth, dividing when cells reach a critical surface area^7^. Nonetheless, our shape growth curve measurements demonstrated that *S. pombe* cells also experience an initial drop in SA/V as they exit stationary phase (Figure S3E). *S. pombe* has a smaller change in cell width (~5%) compared to all bacterial species we imaged, presumably due to its thicker cell wall^8^. The SA/V dynamics in *S. pombe* exhibit more fluctuations and deviate more from our model fitting than most bacterial species, suggesting that other mechanisms may also control *S. pombe* cell shape; nonetheless, both width and length changed along the growth curve in a manner similar to *E. coli*. Taken together, the trends of SA/V are conserved across many microbial species grown in nutrient-rich media and do not dependent on factors such as cell-cycle control, cell size regulation, or cell-wall compositions.

Bacterial cell size during steady-state growth is thought to be relatively constant as growth rate changes across temperatures^9^. Nonetheless, we found that stationary phase *E. coli* cells at 30, 37, and 42 °C had different cellular dimensions and SA/V values. Shape growth curves at 30 °C (Figure S3F) and 42 °C (Figure S3G) revealed initial decreases in SA/V, with approximately the same fractional decrease at minimum SA/V at all three temperatures despite the different times at which the minimum was reached. Interestingly, at 30 °C, a fit of our time-delay model produced ∆*t* = 19 min, ~70% longer than at 37 °C or 42 °C. This difference in ∆*t* is likely due to the slower growth rate at 30 °C, which is ~60% slower during exponential growth compared to 37 or 42 °C: slower growth rates lead to slower protein turnover, and therefore longer time is required for proteome re-allocation.

**SA/V regulation is dependent on nutrient conditions**

We asked whether medium composition affects SA/V dynamics, considering that steady-state cell shape is nutrient-dependent^10^. In the defined medium M9+0.4% glucose, maximum growth rate is ~30% of that in LB, and cells are overall smaller^10^. Shape growth curves in M9 glucose showed that SA/V remained largely constant (<1.5% total relative variation across all time points, defined as standard deviation/mean) (Figure S4A), and similar trends were observed with Davis Minimal (DM) medium+0.4% glucose (Figure S4B), another minimal defined medium containing different salts to M9. In these experiments, while cell length kept increasing for the first 2-3 h, cell width dropped during that period after a brief increase in the first 15-30 min, compensating for the changes in length such that constant SA/V was maintained (Figure S1B, S4C). The initial increase of width in the first 15-30 min post dilution was confirmed by single-cell time-lapse data (Figure S4D). We fit our time-delay model to the shape growth curve in M9, and obtained ∆*t* = 0 min.

We further tested other carbon sources that support slower growth compared to glucose in M9-based media. In all four cases (glycerol, sorbitol, mannitol, and acetate), SA/V also largely remained unchanged with relative variation <2.5% (Figure S4E), and the drop in mean cell width coincided with cell length increase. Similarly, in all four carbon sources, the best fit of our time-delay model gave ∆*t* = 0 min. We postulate that the lack of a time delay for slower growth conditions is because cells require less proteome re-allocation when transitioning from stationary phase to log phase.

We hypothesized that more complex media would require changes in proteome composition, leading to a non-zero ∆*t*. To test this hypothesis, we grew cells in M9 glucose with the addition of six amino acids, which permits slightly higher growth rates (~45% of growth rate in LB)^10^. In this case, the relative variation in SA/V increased slightly (Figure S4F) and the time delay was ∆*t* = 5 min, although the length and width dynamics largely mimicked those in M9 glucose. M9 glucose with casamino acids allowed even faster growth (~70% of growth rate in LB)^10^ and restored LB-like SA/V (Figure S4G) and width changes with ∆*t* = 19 min. In the defined rich medium EZ-RDM, which supports similar growth rates as LB, the dynamics of SA/V and cellular dimensions resembled those in LB (Figure S4H).

**Lipid synthesis does not determine surface area synthesis rate**

We further asked whether the lack of SA/V changes when tuning fatty acid synthesis is caused by the presence of the cell wall, by quantifying the SA/V dynamics in cell wall-deficient spheroplasts^11^. Under treatment of the β-lactam cefsulodin, *E. coli* cells form spheroplasts that stably propagate in osmo-protected media^12^. We diluted overnight spheroplast cultures into fresh media with or without cefsulodin, and quantified cell shape during growth. Due to the irregularity in cell shape of spheroplasts, we used contour length to area ratio (L/A) in the imaging plane as a proxy for SA/V. For rod-shaped cells grown in normal conditions, L/A exhibited a trend similar to SA/V, immediately dropping after growth resumption (Figure S5I), suggesting that L/A is a good proxy for SA/V. In the culture with cefsulodin, cells remained wall-less, and we observed a slight increase in L/A during the first 3 h of growth instead of a decrease. By contrast, while the culture without cefsulodin also exhibited an initial increase in L/A, after ~150 min its L/A decreased to lower than that of the cefsulodin-treated cells (Figure S5I). Interestingly, 150 min is approximately the time scale over which spheroplasts start reverting back to rod-like shape with substantially rebuilt cell walls^12^, indicating that the reduction in L/A is mediated by cell wall synthesis. Thus, these results indicate that envelope growth is likely limited by cell wall rather than membrane synthesis.

To resolve the contrasting behaviors when inhibiting lipid and peptidoglycan synthesis, we asked whether the biosynthesis proteins for lipids and peptidoglycan are differentially regulated. We analyzed a previously published dataset examining *E. coli* proteome dynamics across conditions with different growth rates^13^. The levels of ribosomal proteins were highly correlated with the levels of lipid synthesis proteins, both of which increased monotonically with growth rate (Figure S5J). By contrast, the levels of peptidoglycan synthesis proteins did not correlate with ribosomal protein levels (Figure S5K). Similar trends were observed in data from another proteome study examining proteomic changes during nutrient shifts^14^ (Figure S5L). Taken together, lipid biosynthesis genes are likely similarly regulated as ribosomal genes and directly linked to growth rate, while cell wall-related genes are the limiting factor in envelope synthesis and determine the surface area synthesis rate *β* during stationary phase outgrowth.

**Supplementary Figure Legends**

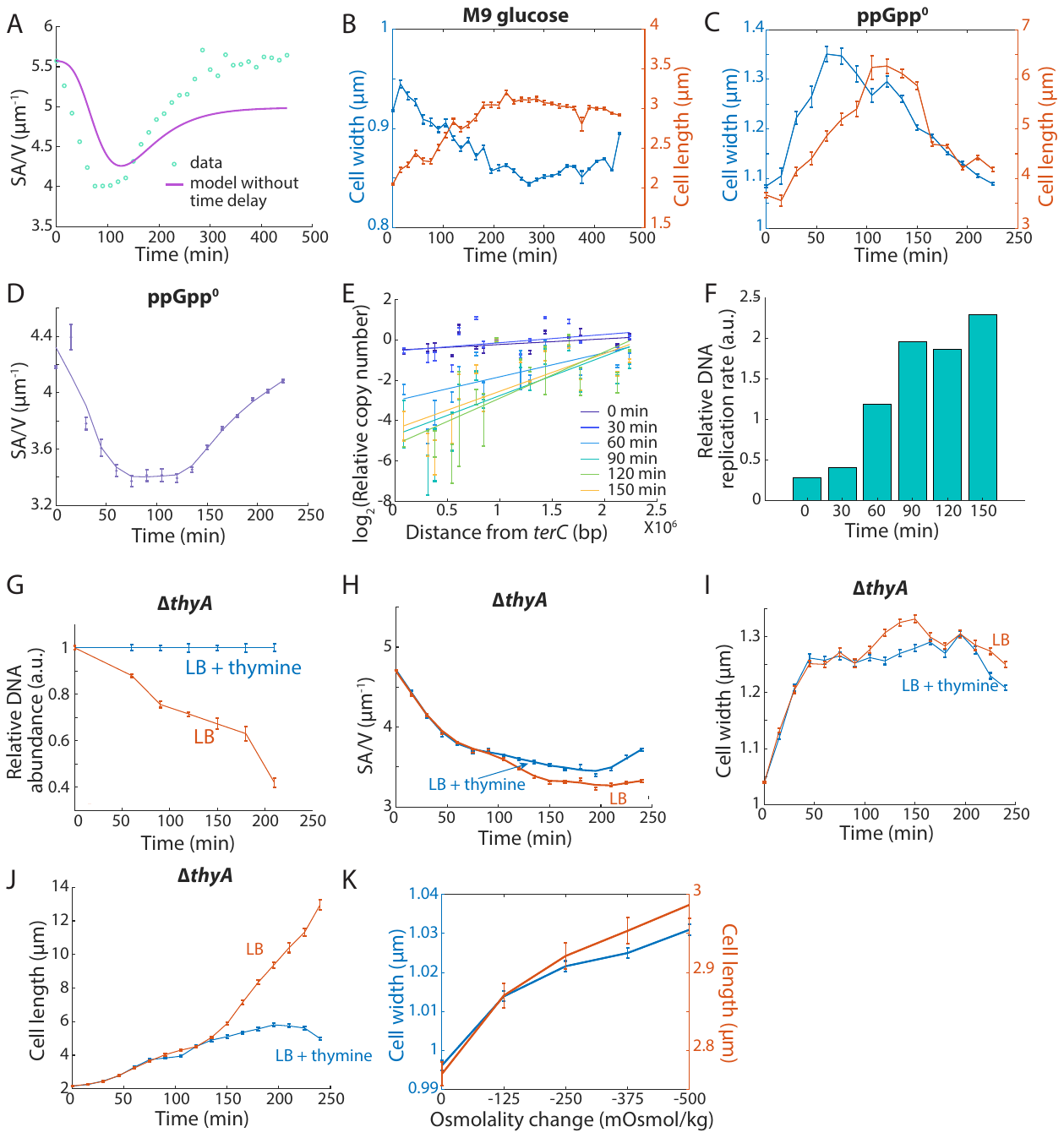

**Figure S1: Cell widening during exit from stationary phase is an active process that optimizes expansion during outgrowth.**

A) The steady-state model (Eq. 3 and 5 in Figure 1E) does not explain the experimentally measured SA/V dynamics.
B) Shape growth curves for cells grown in M9 glucose. Cell width changes in M9 glucose were more complex than in LB. Data points are mean ± s.e.m. with *n* > 200 cells.

C-D) Shape growth curves of an *E. coli* ppGpp^0^ strain. The ppGpp^0^ strain is wider and longer than MG1655, but nevertheless exhibited qualitatively similar cellular dimension dynamics as the wild-type strain, suggesting that SA/V dynamics are not regulated by the stringent response. Data points are mean ± s.e.m., with *n* > 200 cells.

E-F) Estimates of DNA replication rates by fitting log_2_(relative copy number) of genomic loci to their distance from *terC* (E). The slope was close to zero in non-replicating cells, and increased in fast-growing conditions. Replication rate was close to zero for 0- and 30-min samples before starting to increase, suggesting that very little DNA replication occurred during the first 30 min of outgrowth (F). Data points are mean ± S.D. with *n* = 3 replicates.

G) Relative DNA abundance in Δ*thyA* cells with or without addition of external thymine during a shape growth curve. Thymine limitation inhibited DNA replication and led to lower DNA abundance. Data points are mean ± s.e.m., with *n* > 200.

H-J) Shape growth curves for Δ*thyA* cells with or without addition of external thymine. The thymine-deficient culture exhibited less DNA replication but a similar shape growth curve for the first 100 min. After 100 min, cell length increased for the thymine-deficient culture, presumably due to division inhibition caused by the block on DNA synthesis. Data points are mean ± s.e.m., with *n* > 200. Solid lines in (H) are smoothed curves as a guide to the eye.

K) Cell width and length changes after large hypoosmotic shocks. Hypoosmotic shocks altered cellular dimensions by < 10% and had slightly larger effects on relative length changes compared with width, and thus cannot explain the large width expansion observed in shape growth curves. Data points are mean ± s.e.m., with *n* > 50 cells.

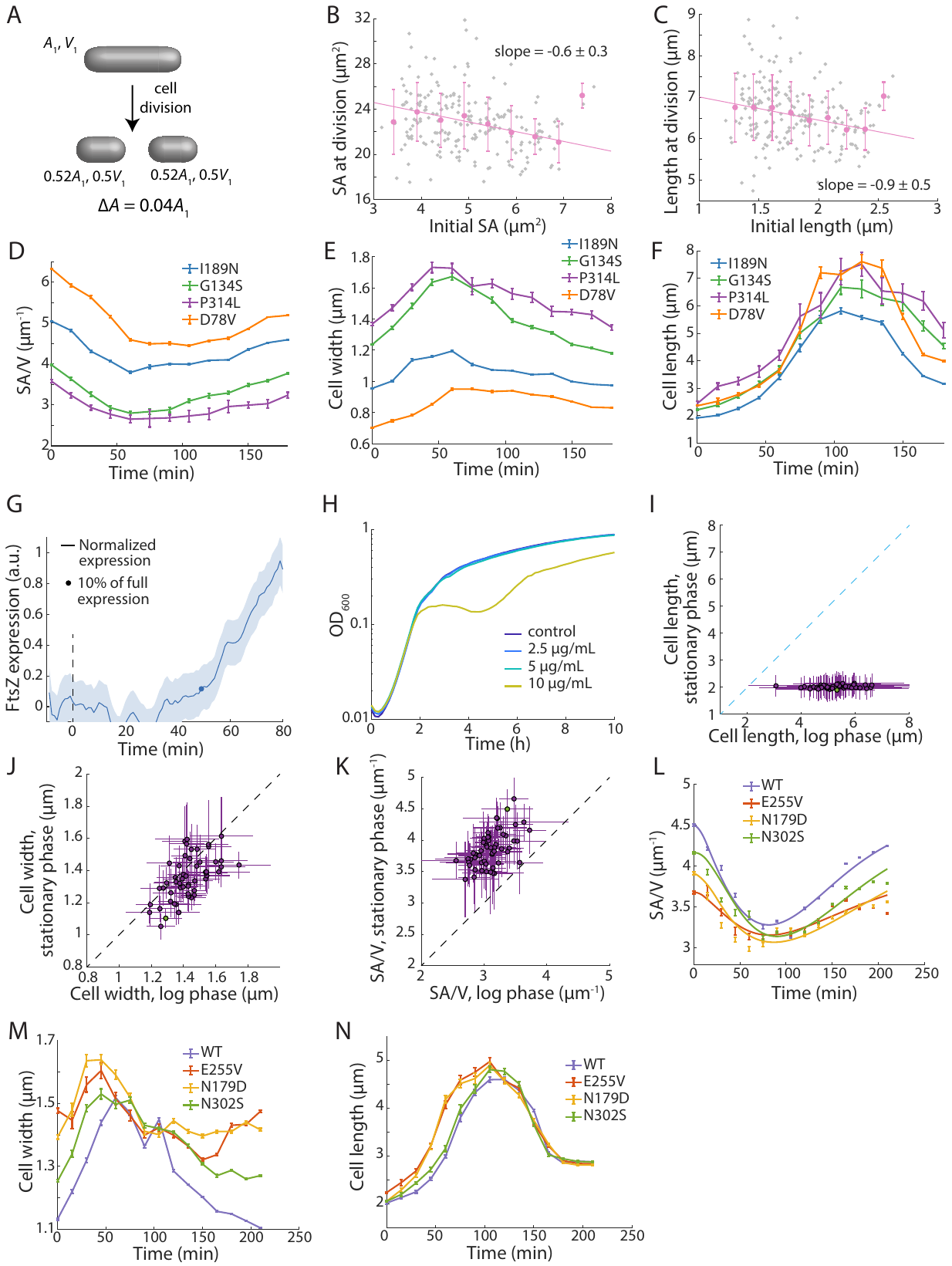

**Figure S2: SA/V dynamics are correlated with cell division and are conserved across cell morphologies.**

A) For a rod-shaped cell starting with surface area *A*_1_ and *V*_1_ (top), cell division adds two hemisphere poles to each of the daughter cells (bottom). The total cell volume remains unchanged, whereas the surface area of each daughter cell is 0.52 *A*_1_. Therefore, cell division requires an additional 4% increase in surface area.

B-C) For cells exiting stationary phase, the surface area (B) and length (C) at division were negatively correlated with the starting surface area or length, indicating deviation from the “sizer” and “adder” models. Gray dots are data points for *n* = 209 individual cells, pink data points are binned mean and S.D. values, which were fit to a linear model.

D) Shape growth curves for MreB cell shape mutants expressing an FtsZsw-mVenus fusion. Despite their quantitatively different SA/V values, all strains exhibited similar trends, with SA/V decreasing from stationary phase and decreased in log phase. Data points are mean ± s.e.m. with n > 200 cells.

E-F) Cell width (E) and length (F) of MreB shape mutants expressing an FtsZ^sw^-mVenus fusion. Despite their quantitatively different widths and lengths, all strains exhibited qualitatively similar width/length dynamics throughout shape growth curves. Data points are mean ± s.e.m. with *n* > 200 cells.

G) FtsZ expression levels were measured using GFP fused to P*_ftsZ_* and normalized from 0 to 1. By contrast with other promoters (Figure 1I), FtsZ levels remained low for ~45 min before increasing, suggesting a delay in up-regulation consistent with the time of first division. The dot represents the time point with 10% increased expression. Data are mean ± standard deviation (S.D.) with *n* > 100 cells.

H) Growth curves of *E. coli* MG1655 in LB with different levels of cephalexin. With 2.5 µg/mL and 5 µg/mL cephalexin, cell growth was not affected; with 10 µg/mL cephalexin, growth rate prior to 2 h remained the same as the untreated control; cell lysis occurs after ~2 h.

I-K) Cell length (I), width (J), and SA/V (K) in stationary (*t* = 0) and log phase (*t* = 120 min) for a library of *E. coli* MreB mutants with different cell sizes. Cell lengths for all strains were very similar in stationary phase (~ 2 µm) but increased by very different extents in log phase. While most strains behaved like MG1655 (Figure 1B,C) with cell width increasing in log phase, a subset had lower width in log phase (J), suggesting more complex width dynamics. Nonetheless, all strains had lower SA/V in log phase compared with stationary phase (K), further validating the generality of SA/V changes. Data points are mean ± standard deviation (S.D.) with *n* > 100 cells. Green dot represents wildtype, and cyan dashed line is *x = y*.

L-N) Shape growth curves for four MreB mutants in (I-K). While the strains had different cell sizes and thus different starting SA/V, the SA/V trends were conserved. For the N255V mutant, cell width dropped to lower than the stationary-phase width by ~120 min, then gradually increased. Such complex changes explain why cell width in log phase is not always larger than that in stationary phase, as shown in Figure S2J. Data points are mean ± s.e.m. with *n* > 200 cells. Solid lines in (L) are fits to the time-delay model, with Δ*t* ~ 15 min for all strains.

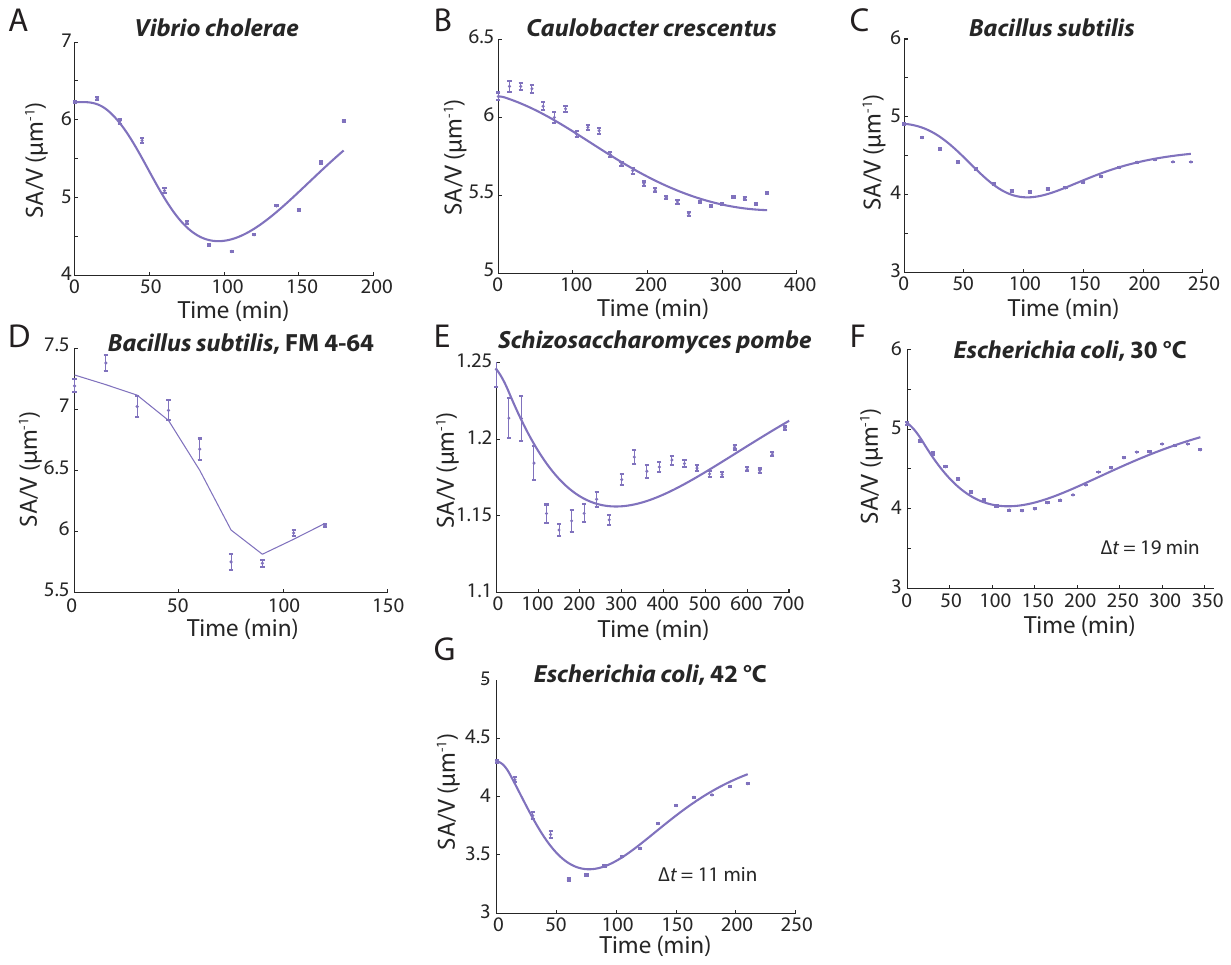

**Figure S3: SA/V dynamics are conserved across species and growth temperatures.**

Shape growth curves for multiple species and temperatures. *V. cholerae* (A) and *C. crescentus* (B) are Gram-negative, *B. subtilis* (C,D) is Gram-positive, and *S. pombe* (fission yeast) is a eukaryotic fungi (E). In all cases, cells exhibited similar trends with SA/V decreasing when cells resumed growth, and then gradually recovering when cell growth slowed down again. Data points are mean ± s.e.m., with *n* > 40 for (E), and *n* > 200 for all other panels. Solid lines are fits to the time-delay model. The time delay at 30 °C (F) was ~70% longer than at 42 °C (G), consistent with the difference in steady-state growth rates.

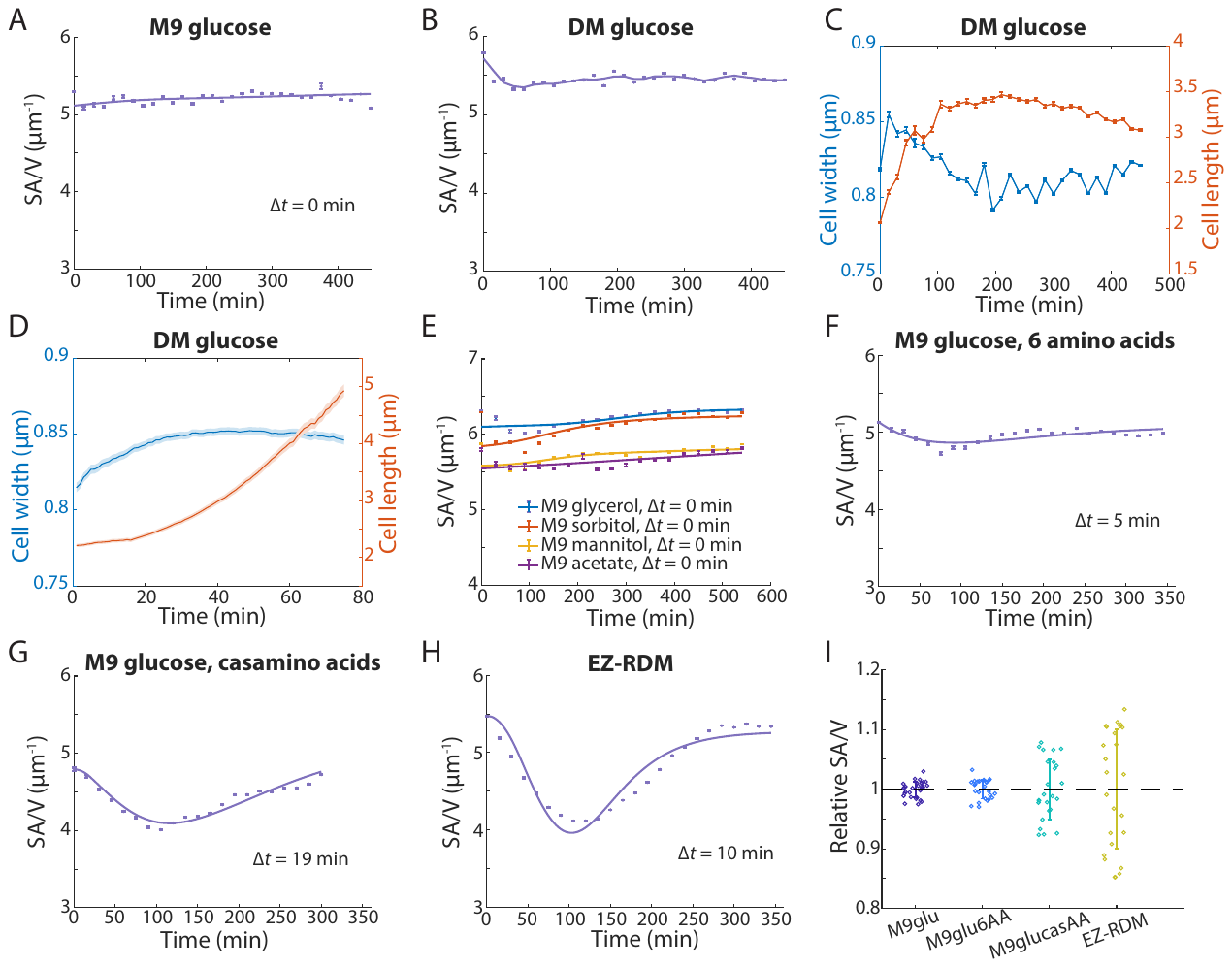

**Figure S4: SA/V regulation is dependent on nutrient conditions.**

A-H) Shape growth curves for MG1655 cells in various media. The range of SA/V values was much smaller for less complex media, particularly those with a single carbon source. Data points are mean ± s.e.m., with *n* > 200. Solid lines in (A) and (E-H) are best fits to the time-delay model.

I) Relative SA/V changes (normalized by mean SA/V across the entire shape growth curve) across media. Richer, more complex media support better growth and result in larger ranges of SA/V. Dots are normalized SA/V of individual time points, and solid lines are mean ± S.D. of the normalized SA/V.

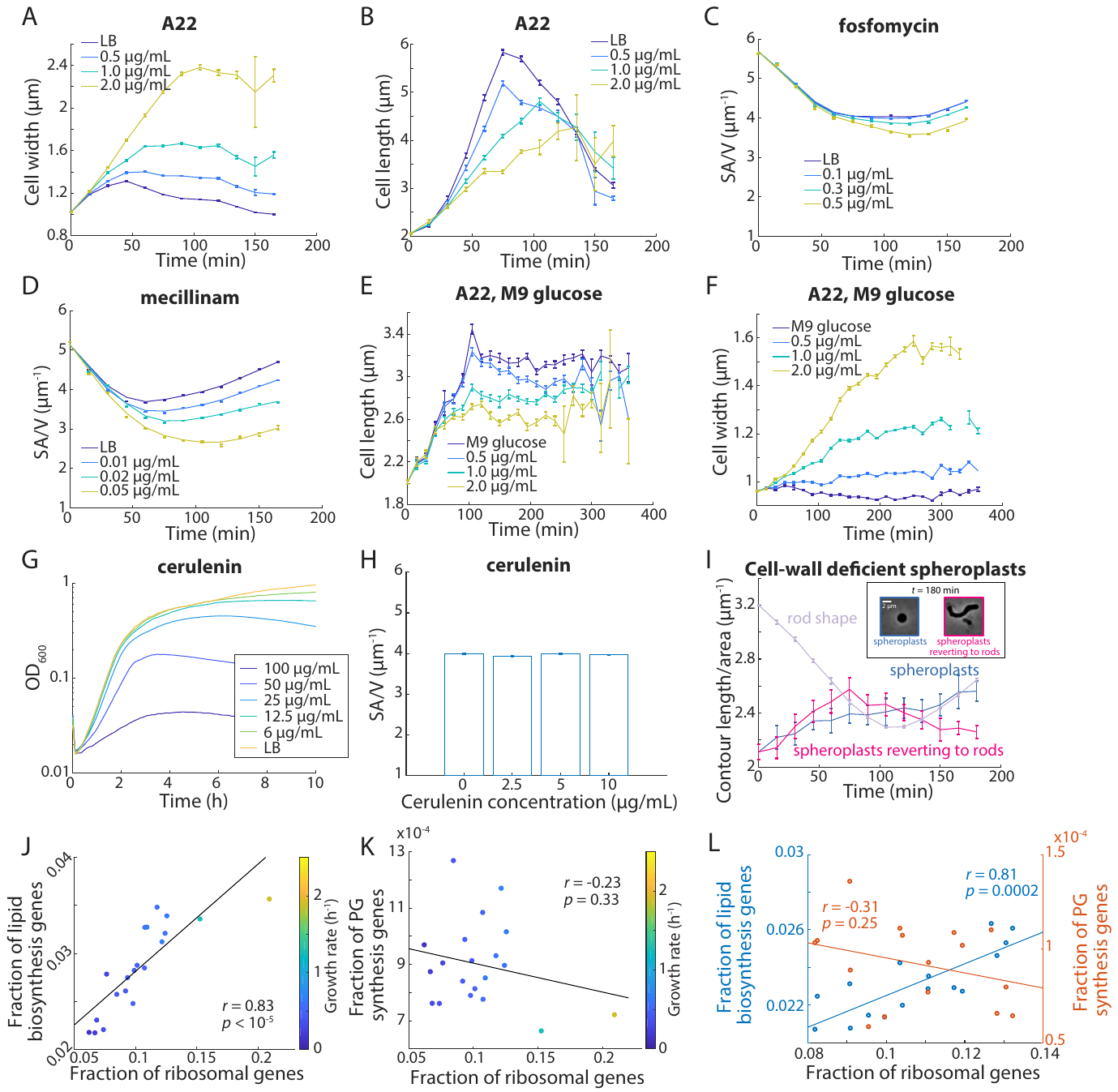

**Figure S5: Inhibiting SA synthesis (*β*) reduces SA/V.**

A-F) Shape growth curves for MG1655 cells treated with A22 (A,B,E,F), fosfomycin (C), or mecillinam (D). In all cases, the antibiotics inhibited cell wall synthesis and therefore reduced SA/V in a dose-dependent manner. Data points are mean ± s.e.m., with *n* > 200 single cells.

G) Growth curves of *E. coli* MG1655 in LB with different levels of cerulenin. Growth rates were largely unaffected with 12.5 µg/mL cerulenin or lower.

H) SA/V for cells after 2 h of cerulenin treatment. Even at the highest concentration tested, cerulenin did not have an observable effect on cell shape. Data points are mean ± s.e.m. with *n* > 1,000 single cells.

I) The ratio between contour length and two-dimensional cell area (L/A) was calculated as a proxy for SA/V in wall-deficient spheroplasts. L/A in rod-shaped cells exhibited a similar trend as SA/V, but spheroplasts increased L/A during growth, suggesting that they are not limited by surface area synthesis. However, for spheroplasts reverting to rod-like shapes, L/A dropped below the non-reverting spheroplasts after ~150 min, when cell-wall regrowth became substantial^12^. Data points are mean ± s.e.m. with *n* > 200 single cells.

J) Proteome data from Ref. ^13^ show that expression levels of lipid biosynthesis genes are highly correlated with ribosomal genes (two-tailed student’s *t*-test), suggesting that lipid levels are directly linked to growth rates. Dots are proteome data in different growth conditions, and the solid line is a linear fit to the data.

K) Proteome data from Ref. ^13^ show that expression levels of peptidoglycan biosynthesis genes are not correlated with ribosomal genes (*p* = 0.33, two-tailed student’s *t*-test), suggesting that by contrast to lipid biosynthesis (J), cell-wall synthesis is regulated differently from genes involved in volumetric growth. Dots are proteome data in different growth conditions, and solid line is the linear best fit of the data.

L) Proteome data from Ref. ^14^ also show that lipid biosynthesis genes are regulated similarly as volumetric growth genes such as ribosomal genes, whereas peptidoglycan synthesis genes are regulated differently.

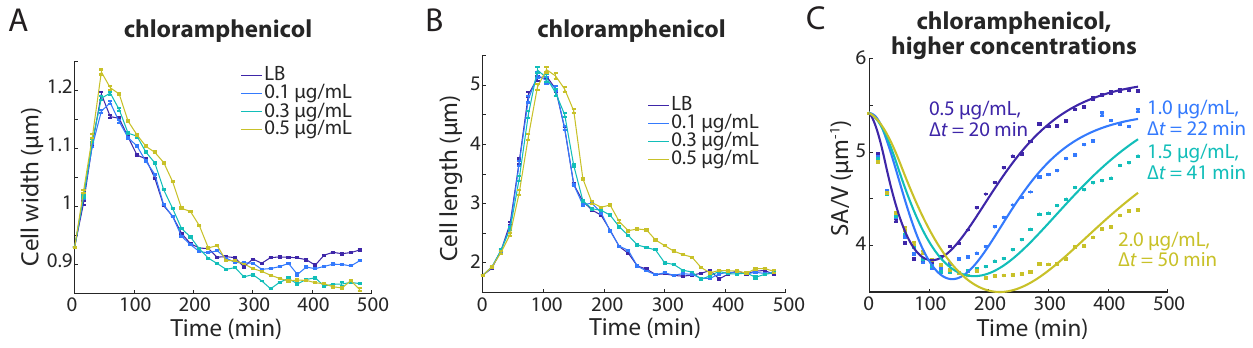

**Figure S6: Inhibiting translation increases the time delay between volume and surface growth.**

A,B) Shape growth curves of MG1655 cells during treatment with low levels of chloramphenicol. The addition of chloramphenicol mostly changed cell width, with larger width in log phase and smaller width when cells re-entered stationary phase. Data points are mean ± s.e.m., with *n* > 200.

C) Higher chloramphenicol concentrations led to even lower SA/V in log phase, and model fitting produced larger ∆*t*. Data points are mean ± s.e.m., with *n* > 200. Smoothed curves are best fits to the model.

**Supplementary Tables**

**Table S1: strains used in this study**

| Strain | Genotype | Source/reference |
| --- | --- | --- |
| *E. coli* MG1655 | *Escherichia coli* K-12 wild-type | CGSC #6300 |
| *E. coli* P*_rplL_*-GFP | BW25113, pSC101, P*_rplL_*-GFP | ^15^ |
| *E. coli* P*_rpsU_*-GFP | BW25113, pSC101, P*_rpsU_*-GFP | ^15^ |
| *E. coli* P*_lacI_*-GFP | BW25113, pSC101, P*_lacI_*-GFP | ^15^ |
| *E. coli* P*_gyrB_*-GFP | BW25113, pSC101, P*_gyrB_*-GFP | ^15^ |
| *E. coli* P*_mrcB_*-GFP | BW25113, pSC101, P*_mrcB_*-GFP | ^15^ |
| *E. coli* P*_murA_*-GFP | BW25113, pSC101, P*_murA_*-GFP | ^15^ |
| *E. coli* P*_ftsZ_*-GFP | BW25113, pSC101, P*_ftsZ_*-GFP | ^15^ |
| *E. coli* ppGpp^0^ | MG1655, ∆*relA*, *spoT::Cm* | ^2^ |
| *B. Subtilis* 168 | *trpC2* | ^16^ |
| *C. crescentus* CB15N | Wild-type | ^17^ |
| *V. cholerae* N16961 | Wild-type | ^18^ |
| *V. cholerae* Δ*wigR* | Δ*wigR* | ^18^ |
| *V. cholerae* pwigR | pHL100 P_lac_-*wigR* | ^18^ |
| *V. cholerae* pwigR-D78E | pHL100 P_lac_-*wigR*-D78E | ^18^ |
| *S. pombe* FC15 | Wild-type 972 | ^7^ |
| *E. coli* MreB-I189N FtsZ-mVenus | MG1655 Δ*mreB* FtsZ-mVenus, pRMmreBCD-I189N | ^5^ |
| *E. coli* MreB-G134S FtsZ-mVenus | MG1655 Δ*mreB* FtsZ-mVenus, pRMmreBCD-G134S | ^5^ |
| *E. coli* MreB-P314L FtsZ-mVenus | MG1655 Δ*mreB* FtsZ-mVenus, pRMmreBCD-P314L | ^5^ |
| *E. coli* MreB-D78V FtsZ-mVenus | MG1655 Δ*mreB* FtsZ-mVenus, pRMmreBCD-D78V | ^5^ |
| *E. coli* MreB-sfGFP | MG1655 Δ*mreB*, pRMmreBCD-msfGFP | ^5^ |
| *E. coli* MreB-sfGFP-E255V | MG1655 Δ*mreB*, pRMmreBCD-E255V-msfGFP | ^5^ |
| *E. coli* MreB-sfGFP-N179D | MG1655 Δ*mreB*, pRMmreBCD-N179D-msfGFP | ^5^ |
| *E. coli* MreB-sfGFP-N302S | MG1655 Δ*mreB*, pRMmreBCD-N302S-msfGFP | ^5^ |
| *E. coli* MG1655 Δ*thyA* | MG1655 Δ*thyA* | ^10^ |

**Table S2: Model fitting results under perturbations of *β* or ∆*t*.**

| Experiment | Condition | *β/β*_0_ | ∆*t* (min) |
| --- | --- | --- | --- |
| LB, A22 | LB | 1.0 | 11 |
|  | 0.5 µg/mL A22 | 0.8 | 11 |
|  | 1.0 µg/mL A22 | 0.7 | 11 |
|  | 2.0 µg/mL A22 | 0.5 | 17 |
| M9, A22 | M9 | 1.0 | 0 |
|  | 0.5 µg/mL A22 | 0.9 | 0 |
|  | 1.0 µg/mL A22 | 0.8 | 1 |
|  | 2.0 µg/mL A22 | 0.6 | 6 |
| LB, chloramphenicol | LB | 1.00 | 11 |
|  | 0.1 µg/mL Cm | 0.99 | 14 |
|  | 0.3 µg/mL Cm | 0.96 | 17 |
|  | 0.5 µg/mL Cm | 0.95 | 19 |
| LB, chloramphenicol, higher concentrations | 0.5 µg/mL Cm | 0.95 | 20 |
|  | 1.0 µg/mL Cm | 0.96 | 22 |
|  | 1.5 µg/mL Cm | 0.97 | 41 |
|  | 2.0 µg/mL Cm | 0.86 | 50 |
